# Sertraline and Carfilzomib Synergize to Target T-cell Malignancies with Serine/Glycine synthesis activity via Cholesterol Dysregulation, Cellular Stress and Immune Modulation

**DOI:** 10.64898/2026.08.17.744660

**Authors:** Paulien Verstraete, Elien Heylen, Anaís Sánchez-Castillo, João Fontela, Lies Matthys, Sofie Meykens, Óscar Herranz, Suraj Verma, Le Minh Thao Doan, Linde Van Aerschot, Jelle Verbeeck, Jonathan Royaert, Michiel Vandenbosch, Ruben Jacobs, Gabriella Dow, Claudio Angione, Annalisa Occhipinti, Daan Dierickx, Jan Cools, Marlies Vanden Bempt, Ilaria Elia, Kim R Kampen, Kim De Keersmaecker

**Author notes:** These authors contributed equally to this work (Paulien Verstraete, Elien Heylen). These authors jointly supervised this work: Kim De Keersmaecker and Kim R. Kampen. CORRESPONDING AUTHORS: Kim De Keersmaecker; Herestraat 49 – ON4, box 810, 3000 Leuven, Belgium;, Kim R. Kampen; Universiteitssingel 50/23, 6229 ER Maastricht, The Netherlands.

## Abstract

**Background:** T-cell acute lymphoblastic leukemia (T-ALL) and peripheral T-cell lymphoma (PTCL) are aggressive hematological malignancies requiring novel therapeutic strategies. The majority of T-ALL and PTCL tumors display metabolic activation and addiction to endogenous serine/glycine synthesis (SSP), providing opportunities for targeted therapy with the clinically used antidepressant sertraline, inhibiting SSP enzymes SHMT1/2. However, sertraline monotherapy only induces cell cycle arrest and has limited efficacy in suppressing disease progression *in vivo*.

**Methods:** Drug synergy of sertraline combined with clinically used proteasome inhibitors carfilzomib and bortezomib was evaluated. Drug effects on cell cycle, proliferation and apoptosis were assessed in T-ALL, PTCL and healthy blood cells using flow cytometry assays. Proteomic, lipidomic and metabolic analyses on drug treated T-ALL cells were performed to elucidate the molecular mechanisms underlying drug synergy, followed by validation of changes of interest, metabolic rescues and shRNA-knockdown of SSP enzymes in T-ALL cells. *In vivo* therapeutic efficacy and immune remodelling were evaluated in an immunocompetent MYCN-overexpressing PTCL mouse model.

**Results:** Sertraline acted synergistically with clinically used proteasome inhibitor carfilzomib to induce cell cycle arrest and apoptosis in T-ALL and PTCL cells with SSP activity, with minimal effects on SSP-inactive T-ALL cells or healthy blood cells. Adding carfilzomib also enhanced the therapeutic efficacy of sertraline in an aggressive MYCN PTCL model. Sertraline rewired cell metabolism towards increased cholesterol uptake and biosynthesis in SSP-active T-ALL cells, and this effect was not obtained by other means of SSP inhibition. In contrast to sertraline, carfilzomib promoted cholesterol efflux. Moreover, carfilzomib reduced total lipid levels, further restricting nutrients in sertraline – carfilzomib treated cells. Additionally, the drug combination impaired mitochondrial respiration and elevated reactive oxygen species (ROS) levels and DNA damage in SSP-active tumor cells, which was rescued by citrate supplementation. Interestingly, these metabolic changes were associated with microenvironmental changes in our mouse model, where the drug combination elevated natural killer T-cells, neutrophils and eosinophils.

**Conclusions:** Our study identifies synergy of sertraline – carfilzomib combination treatment mediated through metabolic impairment and is associated with remodelling of the immune microenvironment. This invites for further clinical investigation of this drug combination as a therapeutic strategy for SSP-active T-cell malignancies.

## BACKGROUND

During cancer development, cancer cells adapt their energy metabolism to supply themselves, and the surrounding niche, with nutrients and biomolecules to facilitate uncontrolled proliferation and survival [1]. For example, cancer cells can activate endogenous serine/glycine synthesis, and even become dependent on this *de novo* serine/glycine synthesis pathway (SSP) to sustain their proliferation. In contrast, most normal cells obtain these non-essential amino acids from their environment and do not produce serine or glycine endogenously. In the SSP, glycolytic intermediate 3-phosphoglycerate (3-PG) is metabolized through three consecutive enzymatic reactions catalyzed by PHGDH, PSAT1, and PSPH to produce serine, which can subsequently be converted into glycine by SHMT (**Figure 1A**). Serine and glycine are important precursors of a variety of biosynthetic pathways. Conversion of serine to glycine generates one-carbon units for the folate cycle, and fuels *de novo* nucleotide synthesis, methylation reactions and antioxidant capacity in cancer cells. In addition, serine serves as a precursor for the synthesis of other amino acids, phospholipids and sphingolipids (**Figure 1A**) [2, 3].

**Figure 1.**
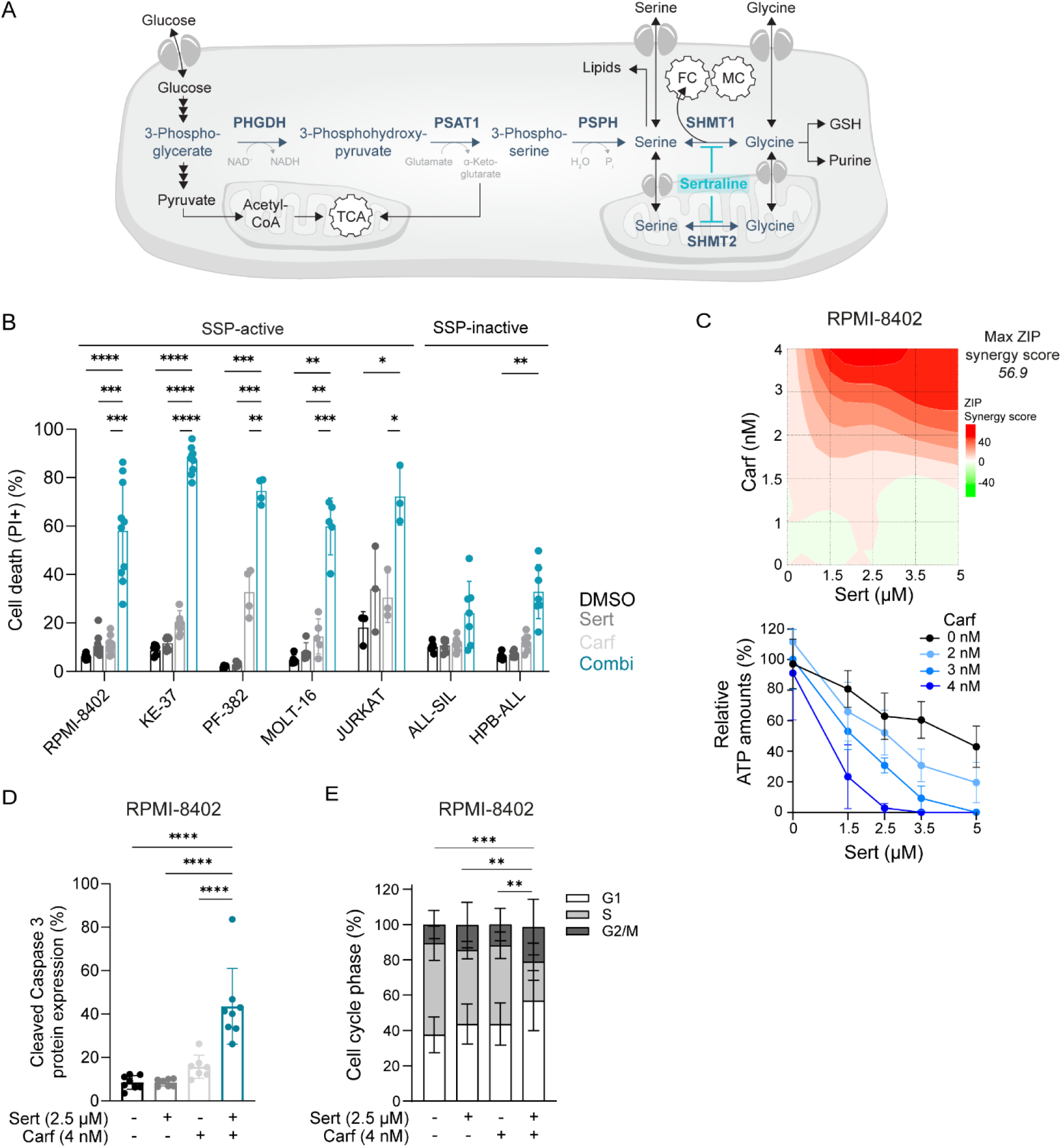
Sertraline acts synergistically with proteasome inhibitor carfilzomib to kill SSP-active T-ALL cell lines. **A)** Scheme of the *de novo* serine/glycine synthesis pathway (SSP). FC = folate cycle; MC = methionine cycle; TCA = tricarboxylic acid cycle. **B)** Cell death analysis measured by propidium iodide (PI) flow cytometry staining of SSP-active (RPMI-8402, KE-37, PF-382, MOLT-16 and JURKAT) and SSP-inactive (ALL-SIL and HPB-ALL) T-ALL cells after 48h of treatment with DMSO, sertraline and/or carfilzomib (n≥3). For each cell line, the drug doses are corresponding to those at which a maximal ZIP synergy score was reached (RPMI-8402: Sert 2.5 µM, Carf 4 nM; KE-37: Sert 5 µM, Carf 1 nM; PF-382: Sert 5 µM, Carf 0.8 nM; MOLT-16; Sert 5 µM, Carf 2 nM; JURKAT: Sert 5 µM, Carf 4 nM; ALL-SIL: Sert 5 µM, Carf 3 nM; HPB-ALL: Sert 5 µM, Carf 3 nM). **C)** Relative ATP levels measured by ATPlite assay and corresponding ZIP-score synergy map of RPMI-8402 treated with sertraline, carfilzomib and their combination therapy for 48h (n≥3). A ZIP synergy score above 10 indicates synergy. **D)** Cell death analysis using cleaved caspase 3 flow cytometry after treatment of RPMI-8402 cells with sertraline and/or carfilzomib for 48h (n≥3). **E)** Quantification of BrdU-PI flow cytometry staining in RPMI-8402 cells treated with sertraline and/or carfilzomib for 48h (n≥7). In B) the combination therapy was compared with DMSO or single agent treated conditions using Two-Way ANOVA with Tukey’s test. In D) combination therapy was compared with either DMSO or single agent treated conditions using a One-Way ANOVA with Šidák’s test. In E) the S-phase contribution was compared between the different groups using Two-Way ANOVA with Tukey’s test. Data are represented as mean ± standard deviation. Individual dots represent independent observations. *P value < 0.05, **P value < 0.01, ***P value < 0.001, ****P value < 0.0001.

Genomic alterations driving upregulation of the SSP have been identified in various cancer subtypes, including blood cancers [4]. Copy number gains of *SHMT2*, encoding the mitochondrial variant of SHMT, occur in B-cell lymphoma [5]. In the same disease, the c-MYC proto-oncogene also transcriptionally upregulates SSP enzyme encoding genes [6]. Furthermore, in acute myeloid leukemia (AML), internal tandem duplications in tyrosine kinase *FLT3* (FLT3-ITD) enhance the SSP via the mTORC1-ATF4 axis [7]. Also, the majority of T-cell acute lymphoblastic leukemia (T-ALL) patient samples and cell lines overexpress the SSP enzymes. Importantly, this overexpression also results in elevated SSP pathway activity, and shRNA knockdown of SPP enzymes impairs T-ALL proliferation and expansion in mice, showing metabolic dependence of these T-ALL cells on the SSP [8]. Several drivers of this SSP metabolic addiction in T-ALL have been identified, including the ribosomal RPL10 R98S mutation that upregulates transcription and translation of PSPH, as well as transcription factor NKX2-1 [4, 8, 9].

Despite advances in treatment options for patients with blood cancer over the past decade, there is still a significant need for novel therapies due to therapy resistance, relapse, severe side effects, and poor survival rates in T-ALL and peripheral T-cell lymphoma (PTCL) [10–14]. These diseases are treated with intensive chemotherapy regimens. While overall survival for T-ALL reaches >80%, lethality occurs due to therapy failure and relapse, and there is long-term drug toxicity [15, 16]. For PTCL, 5-year survival rates are only 10% to 30%, indicating the need for novel therapeutic strategies [14].

SSP enzymes are only minimally expressed in normal bone marrow and thymus [8], making the SSP an appealing therapeutic target in blood cancers with SSP activity. As such, T-ALL, Burkitt’s and diffuse large B-cell lymphoma (DLBCL) cells are highly sensitive to pharmacological SHMT and PHGDH inhibition [17–19]. Several inhibitors of the SSP enzymes have been developed, but their clinical potential is limited by poor pharmacokinetic properties [2]. However, the clinically used antidepressant sertraline has been shown to act as an SHMT inhibitor [20]. Indeed, sertraline selectively inhibits the growth of cancer cells with metabolic activity of the SSP pathway; inhibits *de novo* serine/glycine synthesis to a similar extent as other established serine/glycine synthesis inhibitors in ^13^C_6_-glucose tracing experiments; binds to SHMT1/2 and inhibits downstream nucleotide synthesis; and it mainly induces cell cycle arrest and not cell death, as observed when inhibiting the SSP by other pharmacological or genetic strategies [20]. In line with this, sertraline shows selective toxicity to SSP activated RPL10 R98S and NKX2-1 T-ALL cells [9, 20]. However, as a monotherapy, the *in vivo* efficacy of sertraline is limited [9, 20, 21].

Here, we demonstrate that sertraline in combination with a proteasome inhibitor synergistically eradicates SSP-active T-ALL and PTCL cells, while having minor effects on peripheral blood mononuclear cells (PBMCs). By applying a multi-omics approach, we found that sertraline and carfilzomib modulate cellular cholesterol metabolism and mitochondrial respiration, causing synergistic induction of cellular stress when combining these drugs. Interestingly, in an aggressive MYCN-driven PTCL mouse model, sertraline – carfilzomib combination therapy also induced infiltration of natural killer T-cells (NKT), neutrophils and eosinophils in the spleen. As sertraline is a clinically used drug, and proteasome inhibitors are in clinical trials for both T-ALL and PTCL, our results open perspectives for novel therapies for these disease entities.

## METHODS

### Cell cultures

Cell lines (**Supplementary Table 1**) were obtained from Deutsche Sammlung von Mikroorganismen und Zellkulturen (DSMZ) or American Type Culture Collection (ATCC). T-ALL cells were cultured in RPMI-1640 (Gibco, 11875093) with 10% Fetal bovine serum (FBS) (Gibco, 10270-106) at 37°C 5%CO_2_ for the initial ATPlite, cell cycle analysis and flow cytometry viability stainings (**Figure 1**). For all following experiments, cells were cultured in serine-glycine-glucose-free RPMI-1640 medium (Teknova, R9660) supplemented with 2.8 mM glucose, with/without 0.3 mM serine and 0.1 mM glycine, with/without 3 or 4 mM citrate (Merck, C0759), with/without cholesterol (Merck, C4951) and with 10% dialyzed FBS (ThermoFisher Scientific, A3382001) or human plasma like medium (HPLM) (Gibco, A4899101) supplemented with 10% dFBS. Cell cultures were tested regularly for mycoplasma infection.

### Ex vivo/in vitro treatments

Cells were seeded at 0.3-2×10^6^ cells/mL and drug treated for 48h or 72h using a D300e Digital Dispenser (Tecan). All compounds used are listed in **Supplementary Table 2**.

### Flow cytometry

Cell viability was determined by staining with Annexin V - Zombie Aqua, propidium iodide (PI) or intracellular anti-cleaved caspase 3. Viable cell counts were determined as the proportion of total cell counts staining negative for Annexin V and Zombie aqua, negative for PI or negative for cleaved caspase 3. Cell cycle analysis was carried out using PI staining together with bromodeoxyuridine (BrdU) incorporation. BrdU incorporation was detected by BrdU antibody staining. ROS levels were determined using CellROX Deep Red flow cytometry staining according to the manufacturer’s protocol. For Nile Red staining, cells were collected and incubated with Nile Red staining according to manufacturer’s protocol. Flow cytometry stainings were measured using a FACS Canto ll (Beckton Dickinson) flow cytometer or MACSQuant X (Miltenyi Biotec) and analysed with FlowJo V10 or MACSQuantify software. For intracellular cleaved caspase 3 and γH2AX stainings, cells were fixed and permeabilized with ice-cold methanol for 30 min. at -20°C. Samples were washed and incubated with a primary and an anti-rabbit secondary antibody. All antibodies and dyes used are in **Supplementary Table 3** and **Supplementary Table 4**.

### ATPlite assay

ATP levels were measured using the ATPlite 1step assay (PerkinElmer, 6016731), measured once at the start of drug incubation (t_0_) and once after 48 hours (t_48_). To calculate relative levels, luminescence values at t_0_ were subtracted from values at t_48_ and normalized to the DMSO control.

### Drug synergy

For assessing synergy of a drug combination, we used SynergyFinderPlus[22] to calculate zero interaction potency (ZIP) scores. ZIP scores above 10 are considered to be synergistic, scores between 0 and 10 indicate additive interactions and scores below 0 point at antagonism.

### T-ALL patient derived xenografts (PDX)

Experiments on human PDX samples were approved and supervised by the UZ Leuven ethical committee (approval S68798). Informed consent was obtained from all subjects. T-ALL PDX samples (XB41 and XB47) that had previously been expanded in NSG mice were cultured in RPMI-1640 medium supplemented with 20% FBS, MEM non-essential amino acids (Gibco, 11140050, 1/100), GlutaMax (Gibco, 35050061, 1/100), sodium pyruvate (Gibco, 11360070, 1/100), 20 ng/mL hIL7 (Peprotech, 200-07), 4 ng/mL hIL2 (Peprotech, 200-02), 20 ng/mL hFLT3L (Miltenyi, 130-093-854) and 25 ng/mL hSCF (Miltenyi, 130-093-991) in 5% CO2 at 37°C.

### PBMC isolation and culture

PBMC collection from healthy volunteers and usage of these PBMCs for drug testing experiments was approved and supervised by the ethics committee research UZ/KU Leuven; approval S68326. Peripheral blood was obtained and collected in EDTA-tubes from healthy volunteers. PBMCs were isolated using SepMate-50 tubes (Stemcell Technologies, 85450) and were cultured in RPMI-1640 supplemented with 20% FBS and 15 mM glutamine (ThermoFisher Scientific, 35-050-061) at 37°C 5% CO2.

### 13C6-glucose tracing total metabolite analysis

RPMI-8402 cells were cultured in serine/glycine-free RPMI-1640 medium (Teknova, R9660) supplemented with 2.8 mM ^12^C_6_-glucose or ^13^C_6_-glucose (Cambridge Isotope Laboratories, CLM-1396-1) and 10% dialyzed FBS (ThermoFisher Scientific, A3382001). After 16h of treatment, 10 µL of cell culture medium was added to 990 µL ice-cold 80% methanol-based extraction buffer. Cells were washed in 0.9% NaCl and resuspended in 300 µL extraction buffer. Extracted metabolites were stored at -80°C for one day, centrifuged and subjected to mass spectrometry analysis by the VIB/KU Leuven metabolomics core facility as described previously [20]. Methanol-based extracted medium and cell samples were further derivatised. In short, medium and cell samples were vacuum-dried overnight at 4°C and dissolved in 20 µL (for media samples) or 10 µL (for cell samples) of 20 mg/mL methoxyamine hydrochloride (MOX)-pyridine (MOX, Merck, 226904; Pyridine, Merck, 270970) solution. Samples were vortexed and incubated at 37°C for 1h30 min. Samples were sent for gas chromatography mass spectrometry analysis to the Laboratory of Metabolic Regulation of Cell function, Department of Cellular and Molecular Medicine, KU Leuven. Samples were then analyzed as previously described [23]. A volume of 10 µL of mouse serum was added to 990 µL ice-cold 80% methanol-based extraction buffer and subjected to mass spectrometry analysis by the VIB/KU Leuven metabolomics core facility as described previously [20].

### *In vivo* drug treatment experiments

Mouse experiments were approved by the KU Leuven animal ethics committee (ECD approval P035/2020) and were executed in accordance with all relevant guidelines and regulations at KU Leuven. A total of 100,000 GFP-positive MYCN PTCL cells [14] were injected in the tail vein in 6-8-week-old C57Bl6/J mice to form lymphomas. Five days after injection, mice were treated with DMSO-vehicle, sertraline (15 mg/kg), carfilzomib (4 mg/kg) or the combination therapy twice a week via intraperitoneal injection (the same drug doses were used for treatments of healthy control mice). To evaluate tumor growth, mice were bled weekly via submandibular punction to determine GFP levels and analyze blood counts using a SCIL-VET (Scil animal care company GmbH) machine. Before sacrifice, plasma samples were collected via submandibular punction. Animals were killed by cervical dislocation when the prior defined humane endpoint was reached, which was defined as displaying any signs of suffering or disease such as ruffled fur and curved posture, neurological symptoms such as hind limb paralysis, reduced moving in the cage or loss of ≥20% of body weight. The humane endpoint was reached after 2 weeks in the experiments we report here. After killing, blood, spleen, thymus and bone marrow samples were collected to determine GFP levels and perform immune panel flow cytometry staining.

### *Ex vivo* culturing MYCN mouse cells

GFP-positive MYCN-PTCL spleen cells were collected from mice and cultured in RPMI-1640 medium supplemented with 20% FBS, MEM non-essential amino acids (Gibco, 11140050, 1/100), GlutaMax (Gibco;35050061, 1/100), sodium pyruvate (Gibco, 11360070, 1/100), 50 µM β-mercaptoethanol (Gibco, 1799301), 100 ng/mL hIL2 (Peprotech, 200-02) and 50 ng/mL hIL7 (Peprotech, 200-07) in 5% CO_2_ at 37°C.

### Flow cytometry immune panel

Single cell suspensions obtained from mouse hematopoietic tissues were washed with PBS, TruStain FcX (Biolegend, 101319) was added and incubated for 10 minutes on ice. Antibodies from 7 different immune panels (**Supplementary Table 3**) were added and incubated for 30 minutes. Cells were washed using FACS buffer (PBS, 1% BSA, 1 mM EDTA and 0.05% NaAzide) and acquired on the MACSQuant X (Miltenyi). Analysis was performed using FlowJo software. Gating strategies can be found in **Supplementary Figure 1** and **Supplementary Figure 2**.

### Lipidomics/proteomics

RPMI-8402 cells were treated with DMSO, sertraline (2.5 µM) and/or carfilzomib (4 nM) for 16h. Cell pellets were washed with 150 mM ammonium bicarbonate and proteomic and lipidomic mass spectrometry was performed by the MultiModal Molecular Imaging Institute (M4I) in Maastricht as previously described [24, 25]. Differential protein expression analysis was performed using the DEP2 R package [26]. Only hits detected in at least 3 replicates (out of 4) per condition were considered. Missing values in the normalized data were imputed using kNN method. Gene set enrichment analysis (GSEA) was performed on ranked genes (sign(log2FC) * -log10(pval)). Lipidomics data were analyzed using LipidSig [27].

### Cholesterol quantification assay

RPMI-8402 cells were treated with DMSO, sertraline (2.5 µM) and/or carfilzomib (4 nM) for 16h. Cell medium samples were collected. Cells were washed with PBS and resuspended in 200 µL Chloroform/MetOH (v2:1) per 1×10^6^ cells. Samples were centrifuged at 14,000 rpm for 10 min. The organic phase was vacuum-dried for 20 min. Cholesterol levels were measured in the collected medium and cell samples using the Amplex Red cholesterol assay kit (ThermoFisher Scientific, A12216) according to manufacturer’s protocol. Measurements obtained from collected medium samples were corrected for basal cholesterol levels in the medium.

### Cholesterol uptake assay

Uptake of extracellular cholesterol was analyzed by fluorometry using NBD cholesterol (600440, Cayman Chemical), a fluorescently tagged cholesterol, according to the manufacturer’s instructions. Briefly, RPMI-8402 cells were treated with either DMSO, sertraline (2.5 µM) and/or carfilzomib (4 nM), in serum-free RPMI-1640 medium, and incubated with NBD cholesterol for 16h. Afterwards, cells were washed twice with PBS, and analyzed by flow cytometry.

### Proteasome Glo

Proteasomal activity in RPMI-8402 cells was assessed using the Chymotrypsin-like, Trypsin-and Caspase-like Proteasome-Glo assay (Promega, G8531) according to manufacturer’s protocol.

### Cell Mito Stress Test

RPMI-8402 cells were treated with DMSO, sertraline (2.5 µM) and/or carfilzomib (4 nM) for 16h. Cells were washed and resuspended in Seahorse RPMI-1640 medium (Agilent, 103576-100) supplemented with XF pyruvate (Agilent, 103578-100, 1/100), XF glucose (Agilent, 103577-100, 1/100) and XF L-glutamine (Agilent, 103579-100, 1/100) and seeded in the Seahorse XF24 cell culture microplates (Agilent, 102340-100). Cells were incubated for 60 min. at 37°C and 0% CO_2_. OCR and ECAR levels were measured over time under basal conditions and upon addition of 1 µM oligomycin (Merck, 75351), 1 µM FCCP (Cayman, 15218-10) and 0.5 µM antimycin A (Merck, A8674).

### Western blotting

An equal number of cells (0.3-1*10^6) were lysed (R&D systems, 895561 or Cell Signaling Technology, 9803) and denatured in 1x Laemmli sample buffer (Bio-Rad, 161-0747) containing 2-mercaptoethanol (Sigma-Aldrich). Proteins were separated on Criterion Tris-Glycine eXtended gels (Bio-Rad) or Novex Tris-Glycine gel (ThermoFisher Scientific) and transferred to PVDF membranes using a Trans-Blot Turbo system (Bio-Rad). The membrane was blocked for 1h in 5% milk powder in TBS-T (TBS with 0.1 % Tween-20). Membranes were incubated with primary and secondary antibodies (**Supplementary Table 5**). Proteins were visualized using chemiluminescence on an Azure C600 (Azure Biosystems). Quantification was performed using ImageJ software.

### shRNA transductions

A pLKO1-mCherry plasmid containing shRNA sequences targeting *PHGDH* or the scrambled control were generated as mentioned before [8]. Glycerol E. coli stocks containing pLV-GFP or mCherry plasmids harbouring shSHMT1, shSHMT2 or their scrambled control were ordered from Vector Builder (**Supplementary Table 6**). Lentiviral transductions were performed by co-transfection of the pLKO1 or pLV plasmids with the packaging plasmid psPAX (Addgene, 12260) and the envelope plasmid VSV-G (Addgene, 14888) into HEK293T cells using JetOptimus transfection reagent (Polyplus, 117-01). RPMI-8402 cells were incubated during 24h with lentiviral supernatant in the presence of 1 µg/mL polybrene (Merck, TR-1003) after which stably transduced cells were sorted on a Sony MA900 instrument to enrich for transduced cells.

### Statistics

Statistical analyses were performed using Prism 10 (GraphPad). Bar plots show the mean and error bars define standard deviation. The number of biological replicates per experiment, the number of experiments performed for each dataset, and the statistical analyses performed are stated in the figure legends. Statistical analyses were performed upon F-test for Equality of Variances. All graphs present individual data points.

## RESULTS

### Sertraline acts synergistically with proteasome inhibition to eradicate SSP-active T-ALL cells, with limited toxicity to normal cells

Because proteasome inhibitors are in clinical trials for T-ALL and PTCL, and the proteasome inhibitor bortezomib acts synergistically with pharmacological or genetic targeting of SSP enzyme PHGDH in multiple myeloma [28, 29], we aimed to assess the combined effect of sertraline with proteasome inhibitors in T-ALL. Before testing these drug combinations, we cultured a panel of T-ALL cell lines under serine-and glycine-deprived media conditions to identify the cell lines with an active SSP (**Supplementary Figure 3**). Cell lines RPMI-8402, KE-37, PF-382, MOLT-16 and Jurkat maintained proliferation and viability in the absence of extracellular serine and glycine and were classified as SSP-active. In contrast, ALL-SIL and HPB-ALL were unable to proliferate under these conditions and were SSP-inactive. Notably, when exposing these cell lines to sertraline combined with proteasome inhibitor carfilzomib, we observed a strong synergistic cytotoxicity in all SSP-active T-ALL cell lines, with maximal ZIP synergy scores up to a value of 64. In contrast, SSP-inactive T-ALL cell lines showed much lower sensitivity towards sertraline – carfilzomib combination therapy (maximal ZIP synergy score values of 13 for ALL-SIL and 21 for HPB-ALL cells) (**Figure 1B-C, Supplementary Figure 4, 6F, 20B**). When treated with the sertraline – carfilzomib combination therapy, SSP-active T-ALL cells showed synergistic induction of apoptotic cell death, as evidenced by caspase-3 activation, and G0/G1 cell cycle arrest, as compared to the monotherapies (**Figure 1D-E, Supplementary Figure 4**). Interestingly, the synergistic induction of apoptosis by sertraline -carfilzomib was consistent across media conditions with different serine/glycine and/or glucose levels (**Supplementary Figure 5**).

To evaluate potential toxicity for non-malignant cells, *in vitro* cultured PBMCs from healthy individuals, which exhibited minimal expression of SSP enzymes (**Figure 2A**), were treated with sertraline, carfilzomib, or the combination, at the concentrations that showed synergy in SSP-active T-ALL cell lines. The sertraline – carfilzomib combination therapy induced 13% increase in cell death in PBMCs, as opposed to 46% in SSP-active RPMI-8402 T-ALL cells, indicating minimal toxicity to healthy cells (**Figure 2B**). Additionally, we tested general toxicity *in vivo* by treating healthy mice with sertraline – carfilzomib combination therapy. Thymocyte development or peripheral white blood cell counts were not affected (**Figure 2C-D**), further emphasizing the specificity of drug effects towards SSP-active tumor cells.

**Figure 2.**
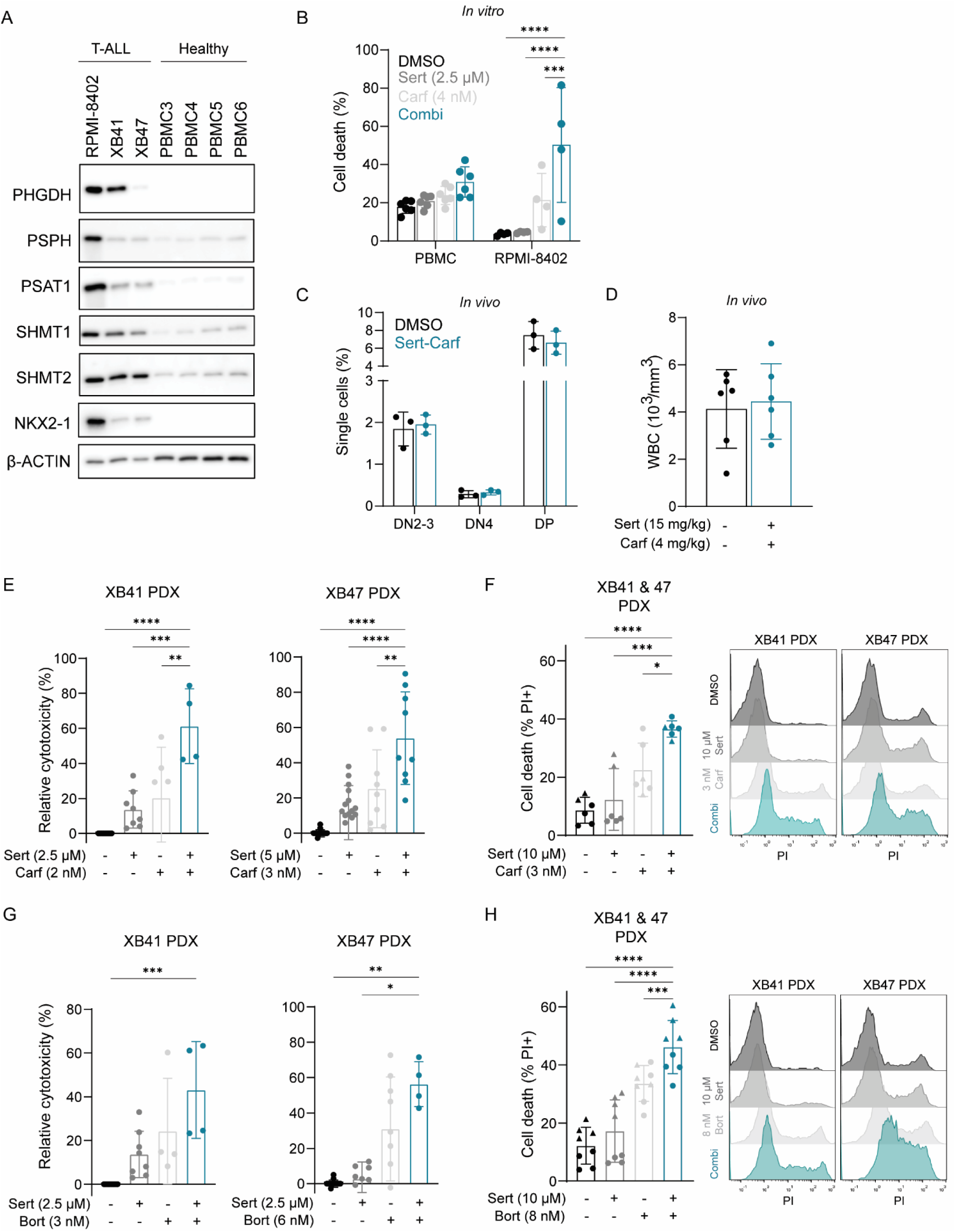
Sertraline synergizes with carfilzomib in T-ALL PDX cells, while having minor impact on normal cells. **A)** Western blot of SSP-driver NKX2-1 and SSP enzymes PHGDH, PSPH, PSAT1 and SHMT1/2 in RPMI-8402 cells, T-ALL PDX cells (XB41 and XB47) and healthy PBMCs. **B)** Cell death in RPMI-8402 cells and PBMCs after 48h treatment with sertraline and/or carfilzomib, measured by propidium iodide (PI) flow cytometry (n≥3). **C)** Thymocyte differentiation stages in healthy mice treated twice weekly with DMSO or sertraline (15 mg/kg) – carfilzomib (4 mg/kg), assessed by flow cytometry (n=3). **D)** White blood cell counts (WBC, 10^3^/mm^3^) in healthy mice treated twice weekly with DMSO or sertraline – carfilzomib (n=6). **E)** Relative cytotoxicity of T-ALL PDX cells (XB41 and XB47) treated *ex vivo* with sertraline, carfilzomib or their combination for 48h, measured by ATPlite (n≥4). **F)** Left: PI-measured cell death in T-ALL PDX cells (XB41: circle; XB47: triangle) treated *ex vivo* with sertraline and/or carfilzomib for 72h (n=6). Right: Representative PI flow cytometry histograms. **G)** Relative cytotoxicity of T-ALL PDX cells (XB41 and XB47) treated *ex vivo* with sertraline, bortezomib or their combination for 48h, measured by ATPlite (n≥4). **H)** Left: PI-measured cell death of T-ALL PDX (XB41: circle; XB47: triangle) cells treated *ex vivo* with sertraline and/or bortezomib for 72h (n=8). Right: Representative PI flow cytometry histograms. In B) combination therapy was compared with DMSO or monotherapies using Two-Way ANOVA with Dunnett’s test. In C) combination therapy was compared with DMSO using Two-Way ANOVA with Šidák’s test. In D) groups were compared using unpaired t-test. In E, F, H) combination therapy was compared with DMSO or monotherapies using One-Way ANOVA with Dunnett’s test. In G) combination therapy was compared with DMSO or monotherapies using a Kruskal-Wallis test (XB41) or One-Way ANOVA (XB47) with Dunnett’s test. Data are represented as mean ± standard deviation. Dots represent independent observations. *P value < 0.05, **P value < 0.01, ***P value < 0.001, ****P value < 0.0001.

We then evaluated whether similar effects could be achieved using the reversible proteasome inhibitor bortezomib. While the sertraline – bortezomib combination induced a synergistic cytotoxic effect in T-ALL cell lines, the obtained maximal ZIP synergy scores were lower than those achieved for sertraline – carfilzomib (e.g. ZIP score of 28 for sertraline – bortezomib versus 57 for sertraline – carfilzomib in RPMI-8402 cells). Furthermore, the association of higher synergy with SSP-activity status of the cells was less consistent for this combination therapy (**Supplementary Figure 6**).

To further assess the therapeutic efficacy of combining sertraline with proteasome inhibitors in clinically relevant models, we used two *ex vivo* cultured T-ALL patient derived xenograft (PDX) samples with SSP activation (XB41 and XB47) (**Figure 2A**). In both T-ALL PDX models, the combination of sertraline and carfilzomib induced higher cytotoxicity and cell death as compared to monotherapies (**Figure 2E-F**). Combining sertraline with bortezomib also enhanced cell death in these T-ALL PDX cells, but synergy was again less pronounced than for the carfilzomib drug combination (**Figure 2G-H**).

In summary, we found that SSP-active T-ALL cells are synergistically targeted by combining sertraline with proteasome inhibitors. The sertraline – carfilzomib drug combination showed the strongest drug synergy on SSP-active T-ALL cells, with limited toxicity to SSP-inactive and healthy cells.

### Sertraline and carfilzomib exert antagonistic effects on lipid and cholesterol homeostasis

To unravel the mechanism of synergy of sertraline – carfilzomib therapy in SSP-active T-ALL cells, quantitative proteomics was performed to compare expression levels of all detectable proteins in RPMI-8402 T-ALL cells treated with DMSO, sertraline and/or carfilzomib. Of >4,000 proteins being detected, 2.1%, 8.9% and 10.5% were significantly differentially expressed as compared to DMSO upon sertraline monotherapy, carfilzomib monotherapy, and sertraline – carfilzomib combination therapy, respectively (**Figure 3A, Supplementary Figure 7**). Not unexpectedly, a large number of proteasomal components were significantly upregulated upon carfilzomib mono-and combination therapy (**Supplementary Figure 8**). Gene set enrichment analysis (GSEA) on the entire proteomics dataset showed significant upregulation of gene sets related to lipid and cholesterol biosynthesis upon sertraline monotherapy, whereas these processes were rather downregulated in the carfilzomib and combination conditions (**Figure 3B, Supplementary Figure 9**). Based on these findings, we performed mass spectrometry-based quantitative lipidomics to assess the effects of our drug treatments on the lipidome. Over 1,200 lipids were identified across all samples. Sertraline treatment altered 2% of lipids covering different lipid classes, whereas carfilzomib monotherapy affected 4.8% of lipid species, with mostly downregulation of phosphatidylcholine (PC) lipids. The combination treatment altered 8.2% of lipid species and, similar to carfilzomib, mainly decreased PCs. In contrast to carfilzomib, the combination treatment also increased lysophosphatidylcholine (LPC) lipids (**Figure 3C-D**). The PC and LPC lipid classes are connected, since LPCs are generated from PCs through phospholipase-mediated fatty acid cleavage, and LPCs can subsequently be reacylated to PCs by lysophospholipid acyltransferases [30]. Overall, this analysis confirmed profound lipidomic changes by our drug treatments. However, cholesterol could not be detected using this lipidomics approach. Since the proteomics also indicated cholesterol synthesis as a pathway induced by sertraline, and because of emerging data on the relevance of this pathway in T-ALL [31–34], we assessed cholesterol synthesis by the mevalonate pathway in more detail. We confirmed significant protein upregulation of cholesterol biosynthesis enzymes HMGCS1 and SQLE in sertraline (mono and combi) treated T-ALL cell lines (**Figure 4A**). Also, expression of the low-density lipoprotein (LDL) receptor (LDLR), responsible for cholesterol uptake, was significantly upregulated upon sertraline treatment, and sertraline increased the cellular uptake of labeled cholesterol (**Figure 4B-C**). Altogether, this was associated with higher intracellular levels of cholesterol (**Figure 4D**). Interestingly, when inhibiting the SSP genetically by shRNA knockdown of PHGDH or SHMT1/2, or pharmacologically with NCT-503 targeting PHGDH or with SHIN1 targeting SHMT1/2, induction of cholesterol synthesis enzymes was not observed (**Figure 4E-F**), and sertraline was still able to induce cholesterol synthesis enzyme HMGCS1 in PHGDH or SHMT1/2 knockdown cells (**Supplementary Figure 10**). These data support that cholesterol synthesis is not directly induced by sertraline mediated SHMT1/2 inhibition. Interestingly, sertraline treatment only induced cholesterol synthesis enzymes in SSP-active T-ALL cell lines (**Figure 4G**), suggesting a crosstalk with the SSP.

**Figure 3.**
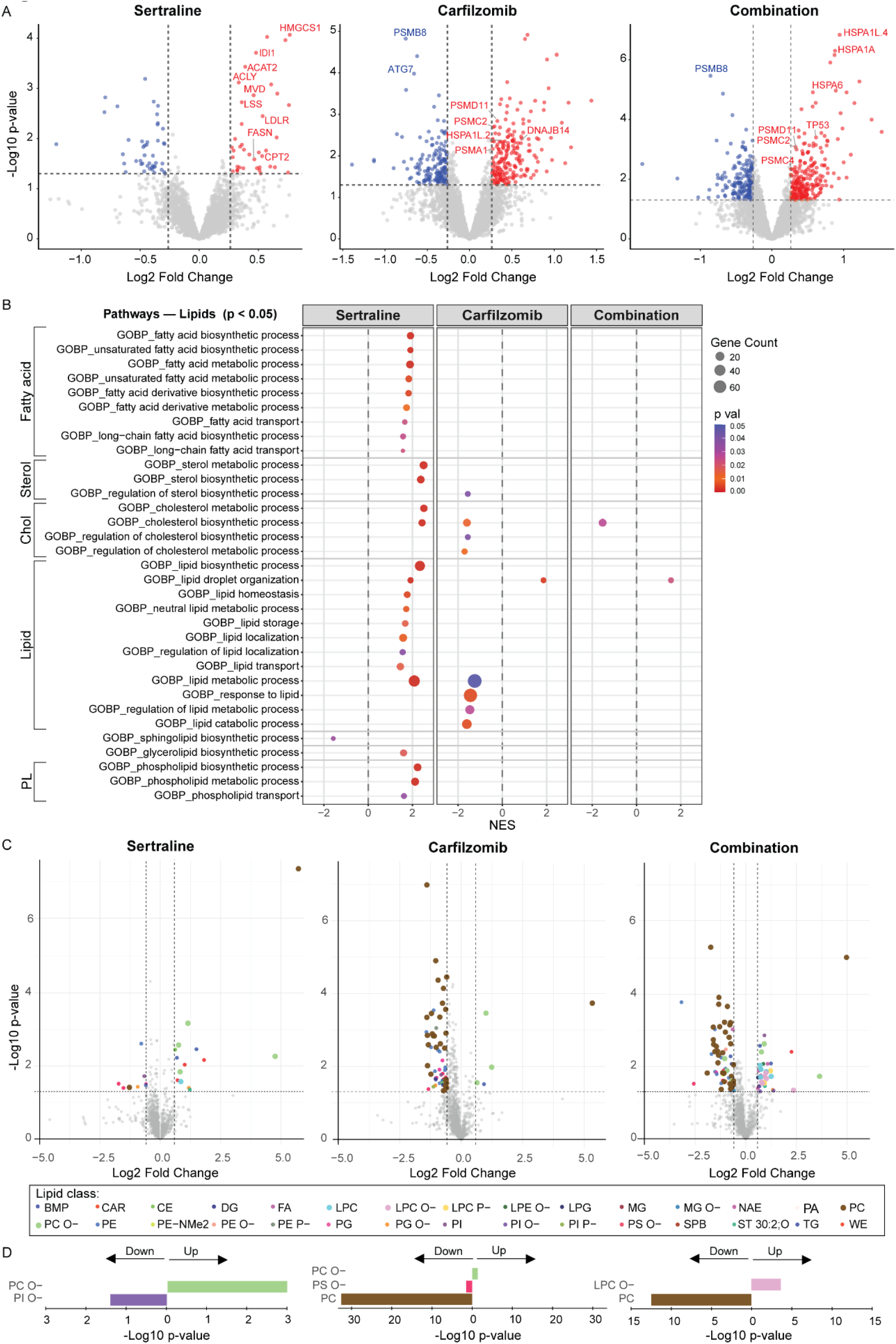
Proteomics and lipidomics analysis of effects of sertraline and carfilzomib monotherapy versus combination therapy. **A)** Volcano plot of proteomics data showing significant proteins altered in RPMI-8402 cells treated with sertraline (2.5 µM), carfilzomib (4 nM) or the combination of both, each time compared to DMSO treated cells. **B)** Gene set enrichment analysis (GSEA) bubble plot of proteomics showing upregulation of proteins related to fatty acid, cholesterol and lipid metabolism in RPMI-8402 cells treated with sertraline monotherapy (2.5 µM), carfilzomib monotherapy (4 nM) or the combination of both compared with DMSO treated cells. All significantly altered pathways from GO-BP related to cholesterol and lipid metabolism with p-value < 0.05 are shown. PL = phospholipid, Chol = cholesterol. **C)** Volcano plot of differentially expressed lipids upon treatment of RPMI-8402 cells with sertraline (2.5 µM), carfilzomib (4 nM) or the combination compared to the DMSO treated condition. p-value < 0.05 and fold change >1.5 were used as cut-off for significance, lipids meeting the significance cut-offs are indicated with colored dots. For the sertraline versus DMSO plot, 1 strongly downregulated lipid (PC O-16:0_22:0, PC O-lipid class, p-value: 3.34E-02 and FC:1.2E-07) could not be plotted as it did not fit on the scaling of the horizontal axis. BMP = Bis(monoacylglycero)phosphate, CAR = carnitine, CE = cholesteryl ester, DG = diacylglycerol, FA = fatty acids, LPC = lysophosphatidylcholine, LPE = lysophosphatidylethanolamine, LPG = lysophosphatidylglycerol, MG = monoacylglycerol, NAE = N-acylethanolamine, PA = diacylglycerophosphates, PC = phosphatidylcholine, PE = phosphatidylethanolamine, PG = phosphatidylglycerol, PI = phosphatidylinositol, PS = phosphatidylserine, SPB = sphingoid bases, ST = sterol lipid, TG = triacylglycerol, WE = wax monoesters. **D)** Enrichment of lipid classes using Over Representation Analysis (ORA). Significant lipid classes are considered when p-value < 0.05.

**Figure 4.**
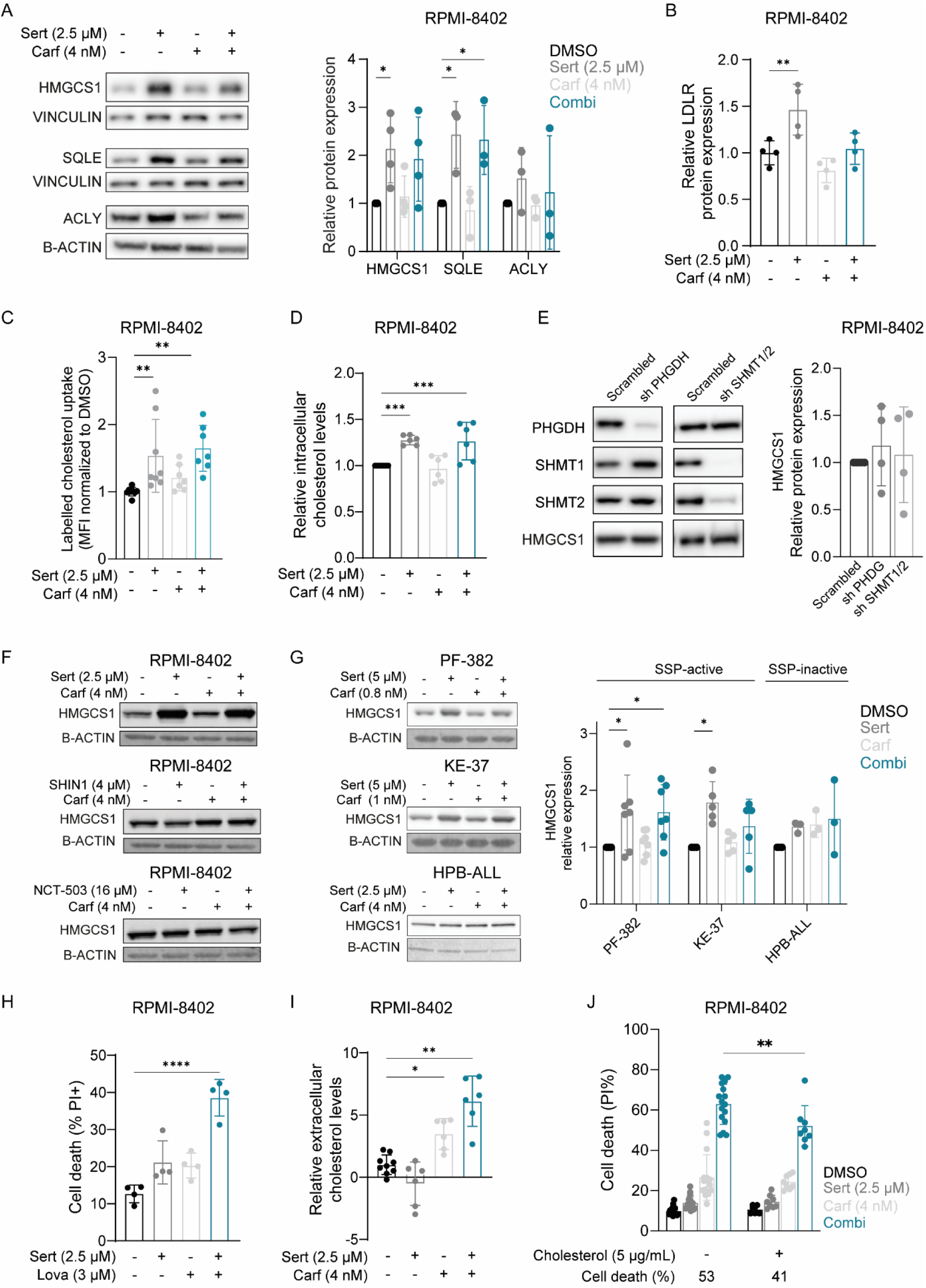
Sertraline – carfilzomib combination therapy modulates cholesterol metabolism. **A)** Western blot (left) and quantification (right) of cholesterol biosynthesis enzymes HMGCS1, SQLE and ACLY in RPMI-8402 cells treated with DMSO, sertraline and/or carfilzomib for 16h (n≥3). **B)** Relative LDLR expression by quantitative proteomics in RPMI-8402 cells treated with DMSO, sertraline and/or carfilzomib for 16h (n≥3). **C)** Relative uptake of fluorescently labeled NBD cholesterol, measured by flow cytometry in RPMI-8402 cells treated with DMSO, sertraline and/or carfilzomib for 16h. **D)** Intracellular cholesterol levels in RPMI-8402 cells treated with DMSO, sertraline and/or carfilzomib for 16h (n≥6). **E)** Western blot of PHGDH, SHMT1, SHMT2 and HMGCS1 in RPMI-8402 cells transduced with scrambled shRNA, PHGDH shRNA or SHMT1/2 shRNA. F) Western blots of HMGCS1 in RPMI-8402 cells treated with DMSO, SSP inhibitor sertraline, SHIN1 or NCT-503; and/or carfilzomib for 16h. **G)** Western blots (left) and quantification (right) of HMGCS1 in SSP-active PF-382, and KE-37 and in SSP-inactive HPB-ALL cell lines treated with DMSO, sertraline and/or carfilzomib for 16h. **H)** Cell death measured by propidium iodide (PI) flow cytometry in RPMI-8402 cells treated with DMSO, sertraline and/or lovastatin for 48h (n=4). **I)** Relative extracellular cholesterol levels in RPMI-8402 cells treated with DMSO, sertraline and/or carfilzomib for 16h (n≥6). **J)** PI-measured cell death of RPMI-8402 cells treated with DMSO, sertraline and/or carfilzomib for 48h with or without 5 µg/mL cholesterol. In A, G) DMSO was compared with monotherapies or combination using Two-Way ANOVA with Dunnett’s test. In B-E, H, I) One-Way ANOVA with Dunnett’s test was used to compare DMSO with sertraline and/or carfilzomib (B-D, H, I), and scrambled shRNA with PHGDH shRNA and SHMT1/2 shRNA (E). In J) treatment effect with and without cholesterol were compared using a Two-Way ANOVA with Šidák’s test. Data are represented as mean ± standard deviation. Dots represent independent observations. *P value < 0.05, **P value < 0.01, ***P value < 0.001, ****P value < 0.0001.

When the cholesterol biosynthesis inhibitor lovastatin was added to T-ALL cells treated with sertraline, cell death was increased (**Figure 4H**), further supporting induction of cholesterol biosynthesis upon sertraline therapy, and even suggesting addiction of sertraline treated SSP dependent T-ALL cells on cholesterol biosynthesis. In contrast to sertraline monotherapy, protein expression of cholesterol biosynthesis enzymes was unaltered upon carfilzomib treatment (**Figure 4A**). However, extracellular cholesterol levels were significantly elevated upon carfilzomib (mono and combi) therapy (**Figure 4I**), whereas cholesterol uptake was unaffected (**Figure 4C**), suggesting more cholesterol efflux. Finally, to assess the role of cholesterol in the anti-leukemic effect of sertraline – carfilzomib combination therapy, we supplemented cholesterol to sertraline – carfilzomib treated RPMI-8402 cells. A 12% rescue in cell death was observed upon cholesterol supplementation (**Figure 4J**), supporting a relevant role of cholesterol metabolism in the anti-leukemic effect of sertraline – carfilzomib treatment. These findings indicate a metabolic antagonism between sertraline and carfilzomib. Whereas sertraline rewires cancer cells towards cholesterol synthesis addiction, carfilzomib limits intracellular lipid availability, in part through enhanced efflux of cholesterol.

### Sertraline – carfilzomib treatment induces citrate depletion, mitochondrial dysfunction and cellular stress responses

Cells synthesize cholesterol and lipids from the metabolite acetyl coenzyme A (AcCoA), which is generated by mitochondrial decarboxylation of glycolytic end product pyruvate (**Figure 5A**). Mitochondrial AcCoA is then converted into citrate, which is exported from the mitochondria to be reconverted into cytosolic AcCoA by ACLY to fuel lipid and cholesterol synthesis. Tracing experiments with ^13^C_6_-glucose showed, in addition to reduced incorporation of glucose-derived carbons in the SSP and downstream nucleotides (**Supplementary 11**), increased incorporation into citrate upon sertraline mono-and combination therapy (**Figure 5B**), potentially fueling cholesterol biosynthesis. To further integrate these ^13^C_6_-glucose tracing with our proteomics datasets, both were used as input to generate an artificial intelligence (AI) metabolic model (**described in Supplementary methods**). This model supported the downregulation of the SSP and upregulation of citrate – cholesterol pathways upon sertraline and combination treatment (**Supplementary Figures 12-14**).

**Figure 5.**
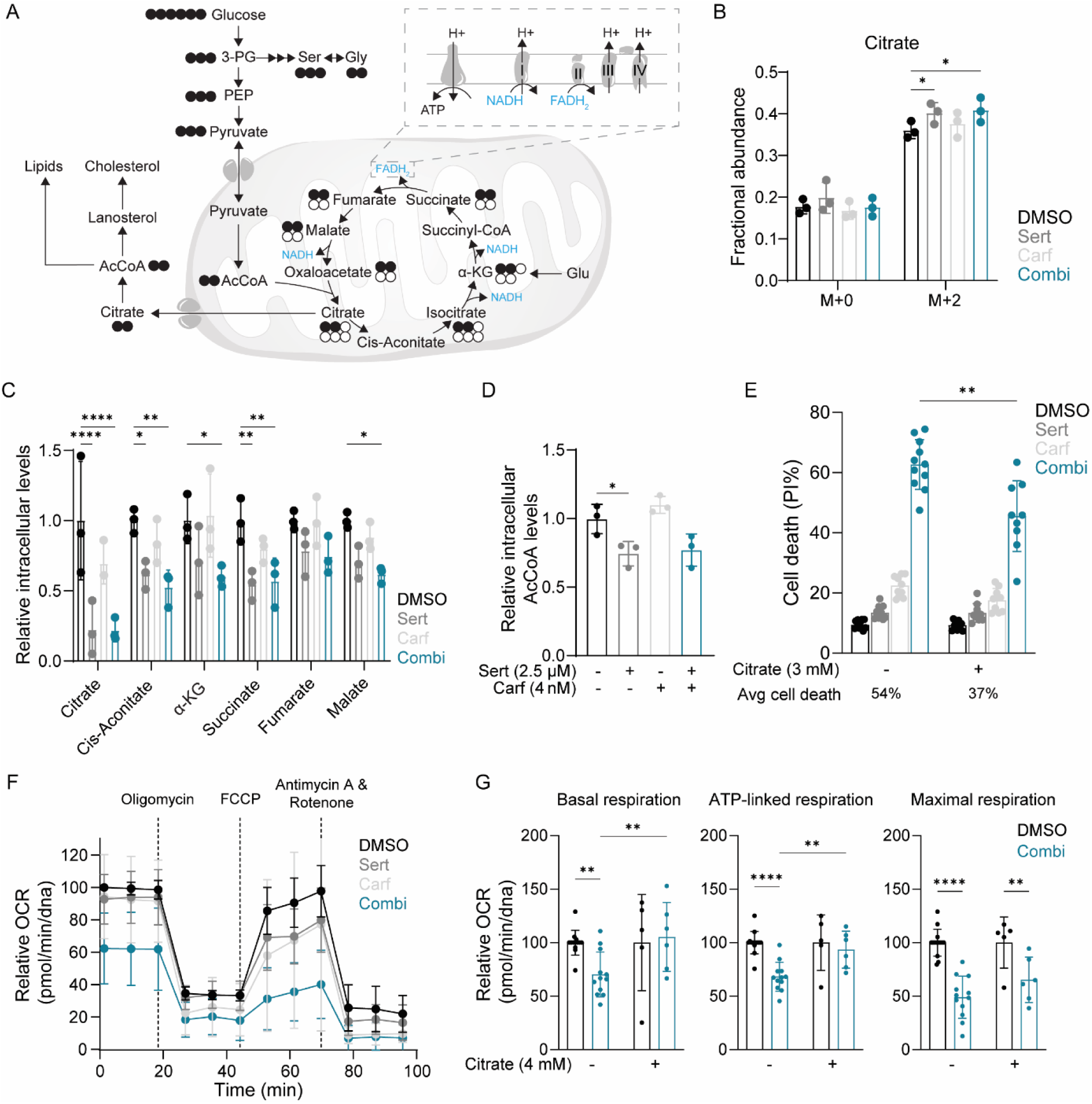
Sertraline – carfilzomib therapy affects mitochondrial metabolism. **A)** Scheme of ^13^C_6_-glucose label distribution into the SSP, TCA cycle and cholesterol/lipid synthesis. Black circles indicate 13C labelled C-atoms, white circles 12C labelled C-atoms. **B)** Fractional abundance of 12-(M+0) and 13C-labeled (M+2) C-atoms in citrate after treatment of RPMI-8402 cells with DMSO, sertraline (2.5 µM) and/or carfilzomib (4 nM) for 16h (n=3). **C)** Relative total intracellular levels of TCA metabolites, normalized for protein input, in RPMI-8402 cells treated with DMSO, sertraline (2.5 µM) and/or carfilzomib (4 nM) for 16h (n=3). α-KG = alpha-ketoglutarate. **D)** Relative total intracellular AcCoA levels, normalized for protein input, in RPMI-8402 cells treated with DMSO, sertraline and/or carfilzomib for 16h (n=3). **E)** Cell death measured by propidium iodide (PI) flow cytometry staining in RPMI-8402 cells treated with DMSO, sertraline (2.5 µM) and/or carfilzomib (4 nM) for 48h with or without 3 mM citrate. **F)** Relative OCR measured using the Seahorse mitostress kit in RPMI-8402 cells treated with DMSO, sertraline (2.5 µM) and/or carfilzomib (4 nM) for 16h. OCR data was normalized for DNA input. **G)** Basal, ATP-linked and maximal respiration in RPMI-8402 cells treated with DMSO or sertraline (2.5 µM) plus carfilzomib (4 nM) and supplemented with or without 4 mM citrate for 16h. In B, C) DMSO was compared to sertraline and/or carfilzomib using Two-Way ANOVA with Dunnett’s test. In D) DMSO was compared to the sertraline and/or carfilzomib using One-Way ANOVA with Dunnett’s test. In E, G) a Two-Way ANOVA with Šidák’s test was used to compare the combination treatment effect with and without citrate (E,G); in (G) combination treatment was also compared with corresponding DMSO in both the absence and presence of citrate. Data are represented as mean ± standard deviation. Dots represent independent observations. *P value < 0.05, **P value < 0.01, ***P value < 0.001, ****P value < 0.0001.

Despite increased incorporation of glucose-derived carbon into citrate (**Figure 5B**), intracellular levels of citrate and AcCoA were significantly reduced in sertraline-and combination-treated cells (**Figure 5C-D**). Given that citrate represents a key metabolic branchpoint between cholesterol biosynthesis and the TCA cycle, depletion of citrate may affect both cholesterol metabolism and mitochondrial carbon availability. Citrate depletion was accompanied by reduced levels of multiple TCA cycle intermediates in sertraline-and combination-treated cells, while the fractional ^13^C_6_-glucose labeling of these metabolites remained largely unchanged (**Supplementary Figure 15**). To determine whether citrate depletion contributed to the observed cytotoxicity, cells were supplemented with exogenous citrate during combination treatment. Citrate supplementation partially rescued cell viability, similar to cholesterol supplementation (**Figure 5E**). Given the central role of the TCA cycle in mitochondrial metabolism, we measured oxygen consumption rate (OCR) and observed that mitochondrial respiration was heavily impaired upon sertraline – carfilzomib combination therapy compared to control or monotherapy (**Figure 5F**). Notably, citrate supplementation improved mitochondrial respiration in sertraline–carfilzomib-treated cells (**Figure 5G**), supporting a contribution of citrate depletion to the mitochondrial phenotype induced by the combination treatment.

ROS levels were increased as well, and the GSH:GSSG ratio reduced when RPMI-8402 cells were treated with sertraline – carfilzomib combination therapy, indicating increased oxidative stress (**Figure 6A, Supplementary Figure 16**). In line with this, γH2AX levels, a mark of DNA damage, were increased when RPMI-8402 cells treated with sertraline – carfilzomib combination therapy (**Figure 6B**). This DNA damage was completely rescued by citrate supplementation (**Figure 6B**). Additionally, RPMI-8402 cells treated with combination therapy showed significantly more lipid droplets (**Figure 6C**), which have been associated with cellular stress responses including lipotoxic stress, endoplasmic reticulum (ER) stress and nutrient deprivation [35, 36]. Consistently, GSEA revealed enrichment of pathways involved in the regulation of cellular responses to stress, and heat shock protein expression was increased in RPMI-8402 cells treated with the combination therapy compared with control or monotherapy (**Figure 6D-E**).

**Figure 6.**
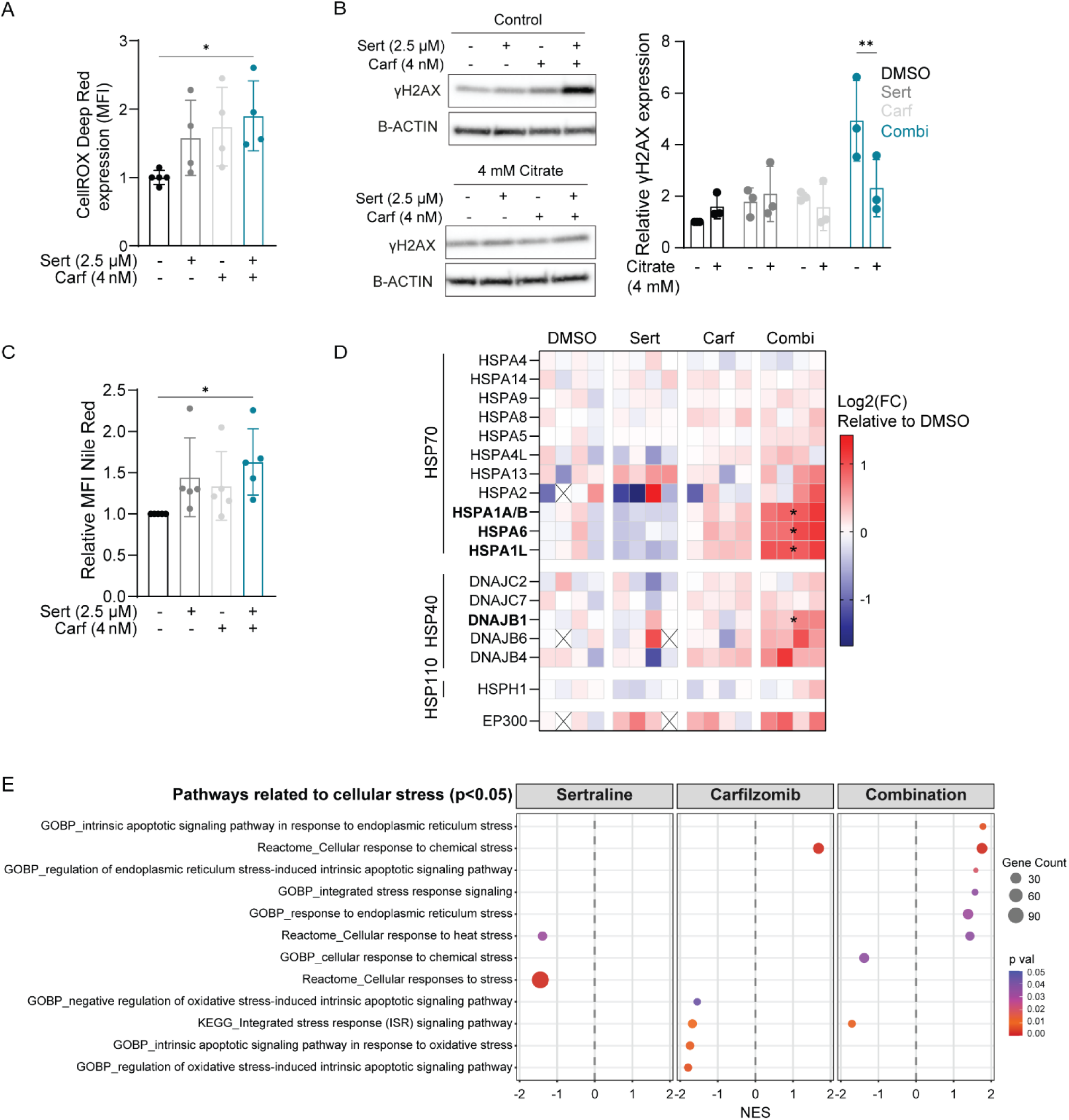
Sertraline – carfilzomib combination treatment induces cellular stress in SSP-active T-ALL cells. **A)** Relative mean fluorescence intensity (MFI) of CellROX ROS flow cytometry staining in RPMI-8402 cells treated with DMSO, sertraline and/or carfilzomib for 16h (n=5). **B)** Representative western blot (left) and the quantification (right) of γH2AX in RPMI-8402 cells treated with DMSO, sertraline and/or carfilzomib and supplemented with or without citrate (4 mM) for 16h (n=3). **C)** Quantification of MFI of Nile Red using flow cytometry in RPMI-8402 cells treated for 16h with DMSO, sertraline and/or carfilzomib (n=5). **D)** Heatmap showing protein expression of heat shock proteins (HSP) in RPMI-8402 cells treated with DMSO, sertraline and/or carfilzomib for 16h (n≥3). **E)** GSEA bubble plot resulting from proteomics of significantly altered proteins (pval <0.05) related to cellular stress in RPMI-8402 cells treated with sertraline and/or carfilzomib compared to DMSO for 16h. In A, C) DMSO was compared to either monotherapies or combination therapy using One-Way ANOVA with Dunnett’s test. In B) for each treatment, the control condition was compared to the citrate condition using Two-Way ANOVA with Dunnett’s test. In D) DMSO was compared to sertraline and/or carfilzomib using Two-Way ANOVA with Holm-Šidák test. Data are represented as mean ± standard deviation. Dots represent independent observations. *P value < 0.05, **P value < 0.01, ***P value < 0.001, ****P value < 0.0001.

Together, these findings support a contribution of citrate depletion, a key metabolic branchpoint between cholesterol biosynthesis and mitochondrial metabolism, to the mitochondrial and oxidative stress phenotype induced by sertraline–carfilzomib treatment. Additional lipid and cellular stress responses may further contribute to the synergistic cytotoxic effect of the combination therapy.

### Sertraline – carfilzomib combination therapy reduces disease burden and modulates the immune landscape in an immunocompetent MYCN-driven T-cell lymphoma mouse model

After having obtained these promising *in vitro* results with sertraline – carfilzomib combination therapy altering cholesterol metabolism, and because of the known role of cholesterol as an immune modulator [37–39], we aimed at establishing an immunocompetent T-cell malignancy model to follow-up on these findings. In PTCL, members of the *MYC*-oncogene family are often overexpressed, which are strong inducers of the SSP [6, 14, 40, 41]. In line with this, MYC-high PTCL patient samples display elevated expression of SSP enzymes and one carbon metabolism genes as compared to MYC-low samples (**Figure 7A-B**). In cells from an immunocompetent MYCN-driven PTCL model [14], we confirmed MYCN binding to active promoter and/or enhancer regions of all SSP enzyme genes. Furthermore, expression of most SSP enzymes was increased in these cells, as well as glycine levels in the blood serum of this MYCN PTCL mouse model (**Figure 7C-F**). In conclusion, similar to its established role in neuroblastoma [41], we show that MYCN is an inducer of SSP expression in PTCL.

**Figure 7.**
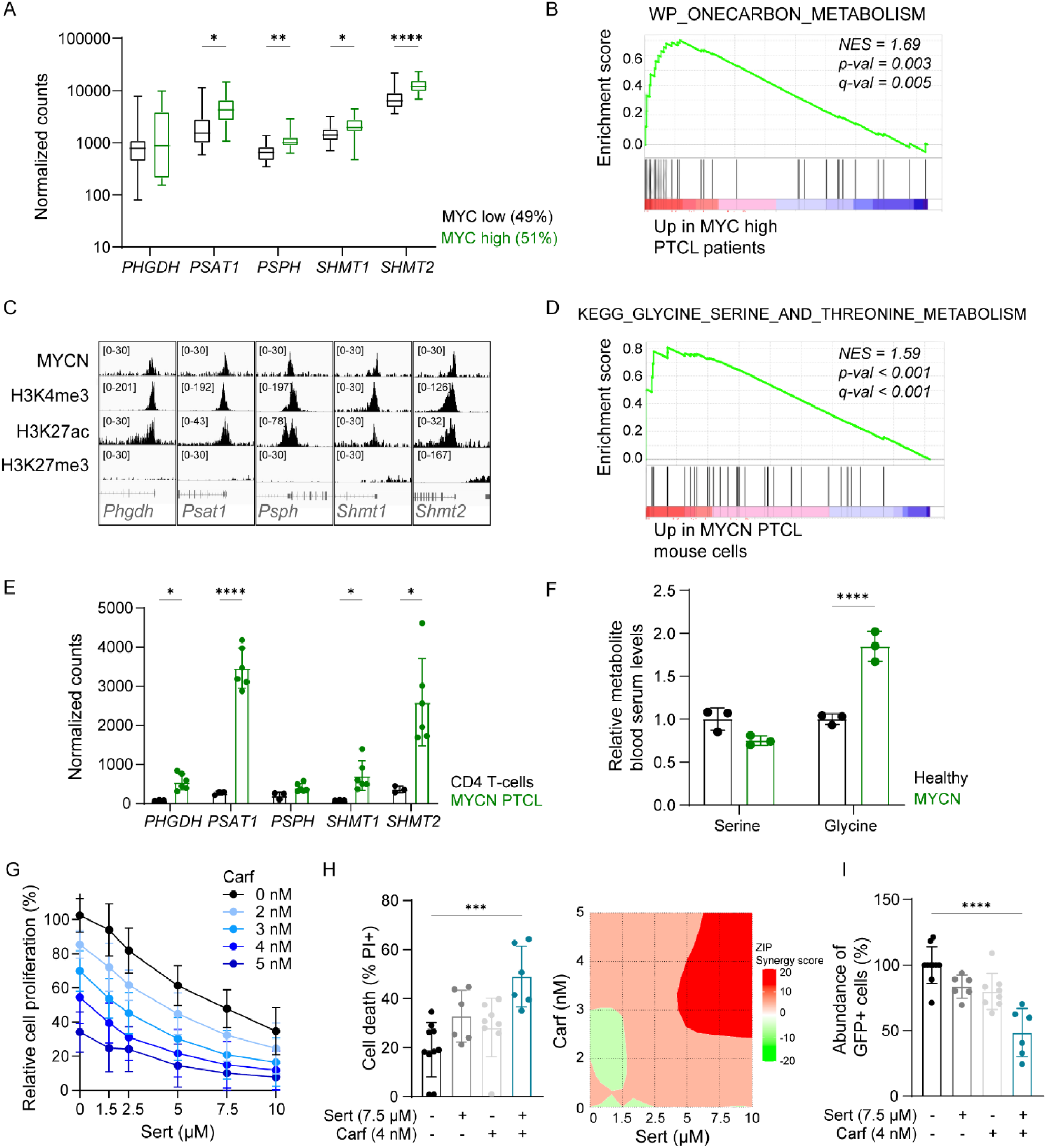
Sertraline – carfilzomib treatment targets MYCN-driven PTCL cells *ex vivo*. **A)** RNA-seq normalized counts of SSP enzyme genes in MYC-high (n=23) and -low (n=22) PTCL patient samples from the published Kyoto and Leuven cohort [14]. **B)** Gene set enrichment analysis (GSEA) of RNA-seq data [14] showing one-carbon metabolism enrichment in differentially expressed genes of PTCL patients with high versus low MYC signature. NES = normalized enrichment score. **C)** MYCN ChIP-seq peaks at promoter (H3K4me3) and enhancer (H3K27ac, H3K4me3-) regions of SSP enzyme genes in MYCN-driven mouse model. Data were downloaded from Gene Expression Omnibus (GEO) (GSE198126). **D)** GSEA of RNA-seq data showing serine metabolism enrichment in differentially expressed genes of CD2 Cre MYCN-driven mouse PTCL cells compared to normal CD4 T-cells. **E)** RNA-seq normalized counts of SSP enzyme genes in CD4+ T-cells or mouse MYCN PTCL cells (n≥3). RNA-seq data for D-E were downloaded from GEO (GSE198126). **F)** Serum serine and glycine abundance in MYCN-driven T-cell lymphoma mice relative to healthy mice (n=3). **G)** Relative proliferation of GFP+ MYCN PTCL mouse cells treated with DMSO, sertraline and/or carfilzomib for 48h. **H)** Cell death measured by propidium iodide (PI) flow cytometry and corresponding ZIP synergy score map of MYCN PTCL cells treated *ex vivo* with DMSO, sertraline and/or carfilzomib for 48h. **I)** Relative abundance of GFP+ MYCN PTCL mouse cells treated with DMSO, sertraline and/or carfilzomib for 48h (n≥6). In A, E, F) a Two-Way ANOVA with Šidák’s test was used to compare SSP gene expression between MYC low/CD4+ T-cells and MYC high/MYCN PTCL (A,E) and serum serine/glycine levels between healthy and MYCN-driven PTCL mice (F). In H, I) cell death and GFP+ cell abundance were compared between DMSO and mono-or combination therapy using Kruskal-Wallis test (H) or One-Way ANOVA with Dunnett’s test (I). Data are represented as mean ± standard deviation. Dots represent independent observations. *P value < 0.05, **P value < 0.01, ***P value < 0.001, ****P value < 0.0001.

To test whether PTCL patients with increased MYCN and serine/glycine enzyme expression may benefit from sertraline, we first tested sertraline – carfilzomib combination therapy on *ex vivo* cultured cells from our MYCN PTCL mouse model [14]. Sertraline and carfilzomib synergistically reduced the percentage of MYCN GFP-positive lymphoma cells, inhibited their proliferation and induced cell death, reaching maximal ZIP synergy scores of 15 (**Figure 7G-I**). We subsequently performed *in vivo* testing in this immunocompetent T-cell lymphoma model [14]. Five days after intravenous injection of the MYCN T-cell lymphoma cells, mice were treated with vehicle (DMSO), sertraline (15 mg/kg), carfilzomib (4 mg/kg) or the combination therapy twice weekly (**Figure 8A**). Due to the aggressiveness of the model, the mice could only be treated 2-3 times before the humane endpoint was reached and all animals were sacrificed on day 13 of the experiment. Nevertheless, a significant reduction in lymphoma disease burden was observed upon combination therapy. Blood samples at the disease end stage showed a 26% reduction of GFP-positive T-cell lymphoma cells, a 17% increased percentage of healthy lymphocytes and reduced white blood cell counts (**Figure 8B, Supplementary Figure 17A**). Additionally, spleen and thymus weight were significantly reduced following sertraline – carfilzomib combination therapy, with a trend towards less GFP expressing lymphoma cells in both organs and in the femurs (**Figure 8C-E, Supplementary Figure 17B**). No toxicity was observed regarding physiological or behavioral signs of pain and/or distress upon combination therapy (**Supplementary Figure 17C**).

**Figure 8.**
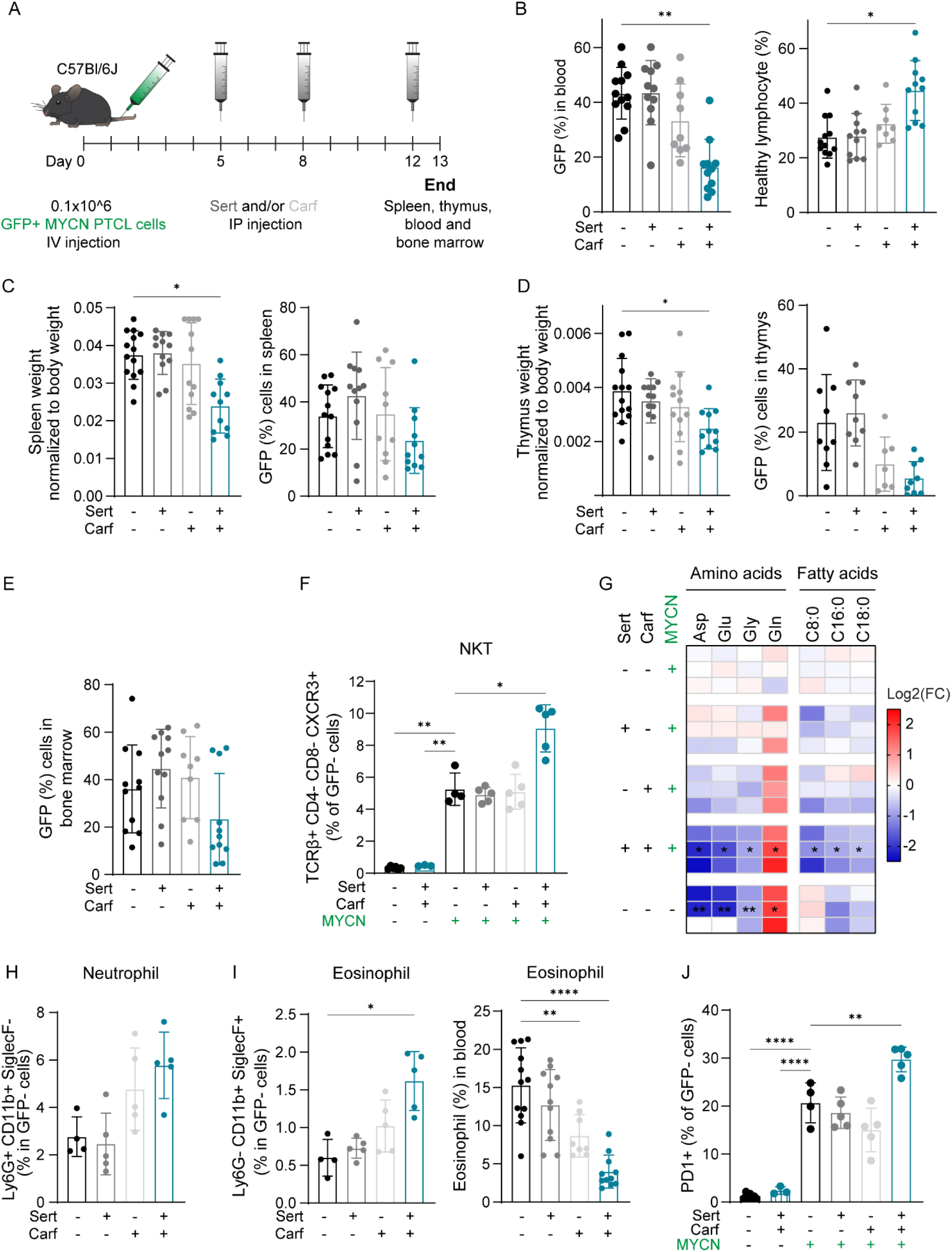
Sertraline – carfilzomib therapy reduces MYCN-driven PTCL disease burden and alters the immune microenvironment *in vivo*. **A)** Schematic overview of the *in vivo* combination treatment experiment. GFP-positive MYCN T-cell lymphoma cells (0.1×10^6) were injected in the tail vein. Treatments with DMSO, sertraline (15 mg/kg, twice weekly), carfilzomib (4 mg/kg, twice weekly) or combination were performed at the indicated time points (day 5, 8 and 12). **B)** Disease burden measured by GFP percentage (left) and healthy lymphocyte percentage (right) in the blood of PTCL mice at sacrifice (day 13). **C-E)** Treatment effects measured at disease end stage (day 13), assessed by spleen weight normalized to body weight and GFP-positive cells in spleen (C), thymus weight normalized to body weight and GFP-positive cells in thymus (D) and bone marrow GFP-positive cells (E). **F)** Effect of sertraline – combination therapy on NKT cells in spleen analyzed by flow cytometry in the GFP-negative population. **G)** Heatmap of relative abundance of amino acids and fatty acids (C8:0 = caprylic acid, C16:0 = palmitic acid, C18:0 = stearic acid) in the serum of MYCN-driven T-cell lymphoma mice treated with DMSO, sertraline and/or carfilzomib and healthy mice. Three biological replicates (mice) are shown per treatment. **H-I)** Effect of sertraline – carfilzomib therapy on percentages of neutrophils (H) and eosinophils (I) in spleen and blood, analyzed by flow cytometry in the GFP-negative population or SCIL-VET. **J)** PD-1 expression in the GFP-negative population of spleen cells of healthy or T-cell lymphoma mice treated with vehicle, sertraline, carfilzomib or the combination. In (G) a Two-Way ANOVA with Dunnett’s test was used to compare mono-or combination therapy to DMSO. The DMSO condition was compared with either healthy mice, single agent treated or combination treated conditions using a One-Way ANOVA with Dunnett’s test (B-F, H-I) or Tukey’s test (J) (n≥3). Data are represented as mean ± standard deviation. Dots represent independent observations. *P value < 0.05, **P value < 0.01, ***P value < 0.001, ****P value < 0.0001.

Because the SSP, sertraline, and cholesterol are known immune modulators [37–39, 42–47], we assessed the effect of sertraline – carfilzomib mono-and combination therapy on immune cell populations and on the expression levels of immune checkpoints in our immunocompetent MYCN T-cell lymphoma model using flow cytometry. Whereas the percentage of CD4^+^ or CD8^+^ T-cells remained the same (**Supplementary Figure 17D**), the natural killer T-cell (NKT) percentage was significantly elevated in the spleens of mice treated with sertraline – carfilzomib combination therapy (**Figure 8F**). This effect was specific to the drug combination, as no changes were observed in healthy animals receiving sertraline or carfilzomib monotherapy. Strikingly, NKT cells were barely detectable in the spleens of healthy mice, even when treated with sertraline – carfilzomib, suggesting a tumor-derived induction of NKT cells upon the combination therapy (**Figure 8F**). Activated type I NKT cells are addicted to glutamine metabolism and take up glutamine from the environment for cytokine production [48]. Interestingly, glutamine levels in the serum of mice treated with sertraline – carfilzomib combination therapy was 3.5-fold increased compared to DMSO (**Figure 8G, Supplementary Table 7**). Moreover, *in vitro* treatment of SSP-dependent T-ALL cells with sertraline – carfilzomib combination therapy increased extracellular levels of glutamine and glutamate (**Supplementary Figure 18**). This indicates that sertraline – carfilzomib treatment increases glutamine levels in the tumor microenvironment, which can attract and activate NKT cells. Moreover, high extracellular levels of palmitic acid (C16:0) inhibit IFNγ and IL-4 production by activated type I NKT cells [49]. Upon sertraline – carfilzomib combination therapy of mice with MYCN PTCL, serum palmitic acid levels were significantly reduced, showing similar levels as healthy mice (**Figure 8G, Supplementary Table 7**). As such, the extracellular glutamine and palmitic acid levels could provide a supportive environment for NKT activation.

Sertraline – carfilzomib combination therapy also increased the percentage of CD11b positive neutrophils and eosinophils in the spleen in our MYCN PTCL model (**Figure 8H-I**). Contrastingly, the percentage of eosinophils in the blood of sertraline – carfilzomib treated mice was reduced, suggesting migration of eosinophils from the blood to the spleen (**Figure 8I**). Cancers with an upregulated SSP have higher expression of the PD-1 immune checkpoint [45]. In accordance, PD-1 expression was increased in spleen cells of MYCN PTCL mice as compared to spleen cells from healthy mice. Strikingly, in the spleen of the lymphoma mice, PD-1 expression was further induced upon sertraline – carfilzomib combination therapy (**Figure 8J**). The expression of other immune checkpoints, such as TIM-3 and TIGIT, was not increased upon sertraline – carfilzomib combination therapy in PTCL mice, and CTLA-4 expression was not detected (**Supplementary Figure 19**).

In summary, sertraline – carfilzomib therapy reduces proliferation and increases cell death of MYCN PTCL cells *ex vivo*. *In vivo* treatment of PTCL-bearing mice with sertraline – carfilzomib combination therapy reduced disease burden in hematopoietic tissues and altered the immune landscape by increasing PD-1 expression as well as the percentage of NKT cells, neutrophils and eosinophils in the spleen.

## DISCUSSION

The SSP is hyperactivated in the majority of T-ALL and PTCL tumors, while normal cells have minimal expression of the SSP enzymes [8, 14]. This difference provides opportunities for therapeutic targeting of the SSP. Moreover, high SSP enzyme expression has been associated with poorer prognosis and drug resistance in various cancer subtypes [3, 18, 28, 29, 50–54], further underscoring the value of the SSP as a therapeutic target.

We show that sertraline, a repurposed antidepressant that inhibits SHMT1/2 [20], has synergistic anti-tumor activity with proteasome inhibitors by inducing cell cycle arrest and apoptosis in SSP-active malignant T-cells. Proteasome inhibitor bortezomib showed less pronounced synergy as compared to carfilzomib, which may be due to the respectively reversible versus irreversible targeting of the proteasome. Interestingly, our -omics analyses and subsequent validations revealed that modulation of cholesterol metabolism is relevant for the drug synergy between sertraline and carfilzomib, as cholesterol supplementation was able to partially rescue apoptosis induction by the combination therapy. More data are emerging that support the critical role of cholesterol homeostasis in hematological malignancies. For instance, cholesterol supplementation has been shown to rescue bortezomib-induced cell death in multiple myeloma cells [55]. Also in T-ALL and T-cell lymphoma, elevated expression and dependence on cholesterol synthesis has been reported, and this can be driven by hypermethylation of N6-methyladenosine in RNA and the HNRNPC-MYC axis, or via Notch-Myc upregulation and increased β-catenin activity [31–34].

We show that treatment of T-ALL cells with sertraline monotherapy increases cholesterol uptake and biosynthesis via the mevalonate pathway, resulting in elevated intracellular cholesterol levels. As this effect on biosynthesis was not observed when inhibiting the SPP pathway genetically or with other established pharmacological SSP inhibitors, cholesterol synthesis is not directly induced by sertraline mediated SHMT1/2 inhibition. Sertraline has previously been shown to promote lysosomal cholesterol accumulation [47, 56], and the induction of cholesterol uptake and synthesis that we see may be a cellular reaction to compensate for this lysosomal lipid accumulation. Such a compensatory induction of cholesterol synthesis also explains why sertraline treated cells are highly sensitized to statins, as we observed. Interestingly, sertraline treatment only induced cholesterol synthesis enzymes in SSP-active T-ALL cell lines, suggesting a crosstalk with the SSP pathway that is of interest to study in the future. We also show that proteasome inhibitor carfilzomib also modulates cholesterol metabolism by increasing cholesterol efflux, and our lipidomics data support that combining sertraline and carfilzomib significantly reduces the intracellular levels of a large number of lipids. The cholesterol efflux effect has previously been observed in macrophages and HEK293T cells, where proteasome inhibition prevented degradation of cholesterol export protein ABCA1/G1 [57]. Our RPMI-8402 metabolic AI model also showed upregulation of ABCA1 (**Supplementary Figure 13**). Cholesterol supplementation could partially rescue the cytotoxicity of sertraline – carfilzomib combination therapy, demonstrating the importance of modulation of cholesterol metabolism for the cytotoxicity of this drug combination. As pharmacological SHMT inhibition by SHIN1 was not inducing cholesterol biosynthesis, it is not surprising that SHIN1 showed lower synergy as compared to sertraline when combined with carfilzomib (**Supplementary Figure 20**). This observation further underscores the importance of the effect of sertraline on cholesterol metabolism in the mode of action of the drug combination and makes sertraline a more effective SSP inhibitor to combine with carfilzomib, as compared to other SHMT inhibitors that do not affect cholesterol metabolism. Furthermore, sertraline is a drug that is already clinically used, whereas other SSP inhibitors have not (yet) made it to the clinic.

Cholesterol and lipids are known immune modulators [58]. Lysosomal cholesterol accumulation upon sertraline treatment has been linked to activation of immunogenic cell death [47, 56], which may be relevant for the immune phenotypes we see in our immunocompetent mouse model. Besides this, tumor derived oxysterols have been shown to attract neutrophils [37, 38]. Other lipids besides cholesterol can also affect the tumor microenvironment. Notably, LPC, which was increased in RPMI-8402 cells treated with sertraline – carfilzomib combination therapy (**Figure 3C-D**), has been reported to stimulate type II NKT cells. In addition, LPC is recognized by the majority of human type I NKT cells [59–61]. We also observed a significant increase in NKT cells in the spleen following the combination treatment *in vivo*. The enhanced presence of neutrophils in our MYCN PTCL model may have been supported by elevated levels of CXCL8 (**Supplementary Figure 21**), a potent neutrophil attracting chemokine that was significantly upregulated in RPMI-8402 cells upon sertraline – carfilzomib combination therapy. Elevated cholesterol in the tumor microenvironment also induces expression of PD-1 and TIM-3 on CD8+ T-cells in lung cancer [39]. We only observed elevated PD-1 expression upon sertraline – carfilzomib treatment of PTCL mice, without increase of other exhaustion markers such as TIM-3. This indicates that PD-1 rather is an activation marker upon sertraline – carfilzomib combination therapy in our MYCN PTCL model. Interestingly, expression of PD-1 on activated iNKT cells has been observed [62], indicating that the increased PD-1 expression upon sertraline – carfilzomib treatment could be explained by the increase of NKT cells in our PTCL model. It is worth noting that the observed changes in the immune microenvironment upon sertraline – carfilzomib therapy may actively contribute to the therapeutic efficacy or may merely be a consequence of tumor regression. To distinguish these scenarios, experiments in immunocompromised mice would be needed. Unfortunately, and in contrast to the C57Bl/6J mice of our PTCL model, NSG mice experienced major discomfort and toxicity of sertraline and carfilzomib monotherapy, preventing us from working in an immunocompromised background.

Serine deprivation and PHGDH inhibition have been shown to impair the TCA cycle in solid tumors [63–65]. In our experiments, despite an equal fractional contribution of glucose-derived carbon to TCA cycle intermediates, total levels of these metabolites were reduced upon sertraline (combination) treatment (**Figure 5C**, **Supplementary Figure 15**). This may be explained by reduced usage of glutamine for the TCA cycle, as observed upon serine deprivation and SSP inhibition in HCT116 cells and NOS1 cells [63, 64]. In line with this, we saw that intracellular glutamate levels were significantly reduced upon sertraline – carfilzomib combination therapy due to enhanced secretion (**Supplementary Figure 18, 22**). However, the reduction in TCA intermediates we saw was larger than the reduction of intracellular glutamate. In this regard, increased diversion of TCA intermediates into other metabolic pathways (cataplerosis) may further contribute to the observed decrease in TCA metabolite levels. Strikingly, although TCA levels were equally reduced by sertraline mono-and combination therapy, mitochondrial respiration was only impaired upon sertraline – carfilzomib combination therapy. This suggests that cells under sertraline monotherapy can replenish the reducing equivalents NADH and/or FADH_2_ that are needed for the electron transport chain, whereas this compensatory mechanism fails when carfilzomib is added. In this regard, fatty acid and fatty acyl CoA metabolism was highly induced upon sertraline monotherapy as shown by increased FASN expression. In contrast, sertraline – carfilzomib combination therapy displayed significantly decreased levels of FASN. Furthermore, we noticed that expression of CPT2, a key enzyme in mitochondrial fatty acid oxidation, was significantly upregulated upon sertraline monotherapy solely (**Supplementary Figure 23**). Because fatty acid oxidation generates FADH_2_ and NADH to fuel the electron transport chain, enhanced fatty acid synthesis and oxidation during sertraline monotherapy may compensate for reduced TCA cycle metabolite pools and thereby maintain mitochondrial respiration. Conversely, sertraline – carfilzomib combination therapy might impair fatty acid synthesis and oxidation, resulting in impaired mitochondrial respiration. In agreement with this, we show that supplementation with citrate, a precursor of AcCoA needed for fatty acid synthesis, rescued mitochondrial respiration and DNA damage.

## CONCLUSION

We identify sertraline combination therapy with proteasome inhibitors as a novel strategy for SSP addicted T-ALL and PTCL cells. By affecting cholesterol and lipid homeostasis, mitochondrial respiration and cellular stress, this combination therapy synergistically induces apoptosis in T-ALL cells and reduces aggressive PTCL disease burden *in vivo*. Furthermore, we show that this combination therapy modulates the tumor immune environment (**Figure 9**). Interestingly, the sertraline mediated induction of cholesterol biosynthesis does not occur when inhibiting the SSP genetically or with other pharmacological inhibitors, which may explain why sertraline shows stronger synergy with carfilzomib as compared to other preclinical SSP inhibitors. Given that sertraline and proteasome inhibitors are clinically used drugs with known clinical pharmacokinetics and toxicities, this study uncovers a novel therapeutic opportunity for T-cell malignancies.

**Figure 9.**
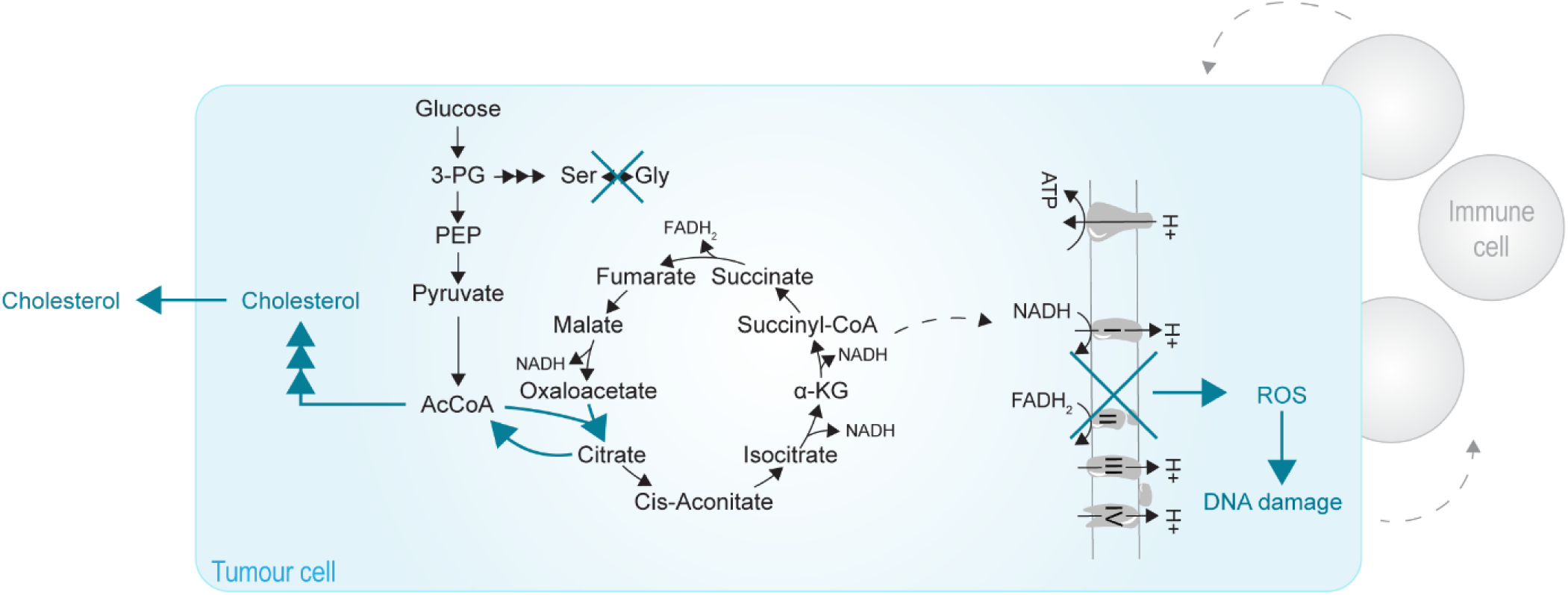
Sertraline – carfilzomib combination therapy disrupts cholesterol and lipid metabolism, mitochondrial respiration, and provokes cellular stress. Scheme of the effects of sertraline – carfilzomib combination therapy (indicated with turquoise color). The figures shows that the combination therapy affects cholesterol biosynthesis and transport and mitochondrial respiration resulting in oxidative stress, DNA damage and cellular stress inside the tumor cell. Moreover, the levels of NKT cells, neutrophils and eosinophils in the immune microenvironment of the tumor cells are elevated by sertraline – carfilzomib combination therapy.

## DECLARATIONS

### Ethics approval and consent to participate

Experiments on human PDX samples were approved and supervised by the UZ Leuven ethical committee (approval S68798). PBMC collection from healthy volunteers and usage of these PBMCs for drug testing experiments was approved and supervised by the ethics committee research UZ/KU Leuven (approval S68326). Mouse experiments described in this study were approved by the KU Leuven animal ethics committee (ECD approval P035/2020) and were executed in accordance with all relevant guidelines and regulations at KU Leuven.

### Availability of data and materials

The data supporting the findings of this study are available from the corresponding author upon reasonable request.

### Authors’ contributions

PV, EH, ASC and JF designed and performed research, analysed the data and wrote the manuscript. LM and SM performed research and analysed data. OH analysed data. MVB generated the T-cell lymphoma mouse model and performed ChIP-seq and RNA-seq. JC reviewed the manuscript and provided PDX samples. DD reviewed the manuscript. LVA performed research. JV and JR provided technical experimental support. SV, LMTD, CA and AO performed AI-based data modelling. MV and RJ performed research and analysed and deposited the proteomics/lipidomics data. GD and IE performed research and analysed the metabolomics data. KRK designed and performed research, analysed data, supervised the study and reviewed the manuscript. KDK designed research, supervised the study, and wrote the manuscript.

## Acknowledgements

We thank the VIB metabolomics core Leuven for performing metabolomics experiments. Lipidomics and proteomics mass spectrometry experiments were performed by the M4I MSI CORE facility at Maastricht University.

## LIST OF ABBREVIATIONS

AcCoA: Acetyl coenzyme A
AI: Artificial intelligence
AML: Acute myeloid leukemia
ATCC: American Type Culture Collection
BrdU: Bromodeoxyuridine
DLBCL: Diffuse large B-cell lymphoma
DSMZ: Deutsche Sammlung von Mikroorganismen und Zellkulturen
ER: Endoplasmic reticulum
FBS: Fetal bovine serum
FLT3-ITD: Internal tandem duplications in tyrosine kinase FLT3
GSEA: Gene set enrichment analysis
HPLM: Human plasma like medium
LDL: Low-density lipoprotein
LDLR: Low-density lipoprotein receptor
LPC: Lysophosphatidylcholine
MOX: Methoxyamine hydrochloride
NKT: Natural killer T-cells
OCR: Oxygen consumption rate
PBMC: Peripheral blood mononuclear cell
PC: Phosphatidylcholine
PDX: Patient derived xenograft
PI: Propidium iodide
PTCL: Peripheral T-cell lymphoma
ROS: Reactive oxygen species
SSP: Serine/glycine synthesis pathway
T-ALL: T-cell acute lymphoblastic leukemia
ZIP: Zero interaction potency
3-PG: 3-phosphoglycerate

