## Supplementary material for "Sertraline and Carfilzomib Synergize to Target T-cell Malignancies with Serine/Glycine synthesis activity via Cholesterol Dysregulation, Cellular Stress and Immune Modulation": Supplentary Figures, Tables and Methods

### SUPPLEMENTARY FIGURES

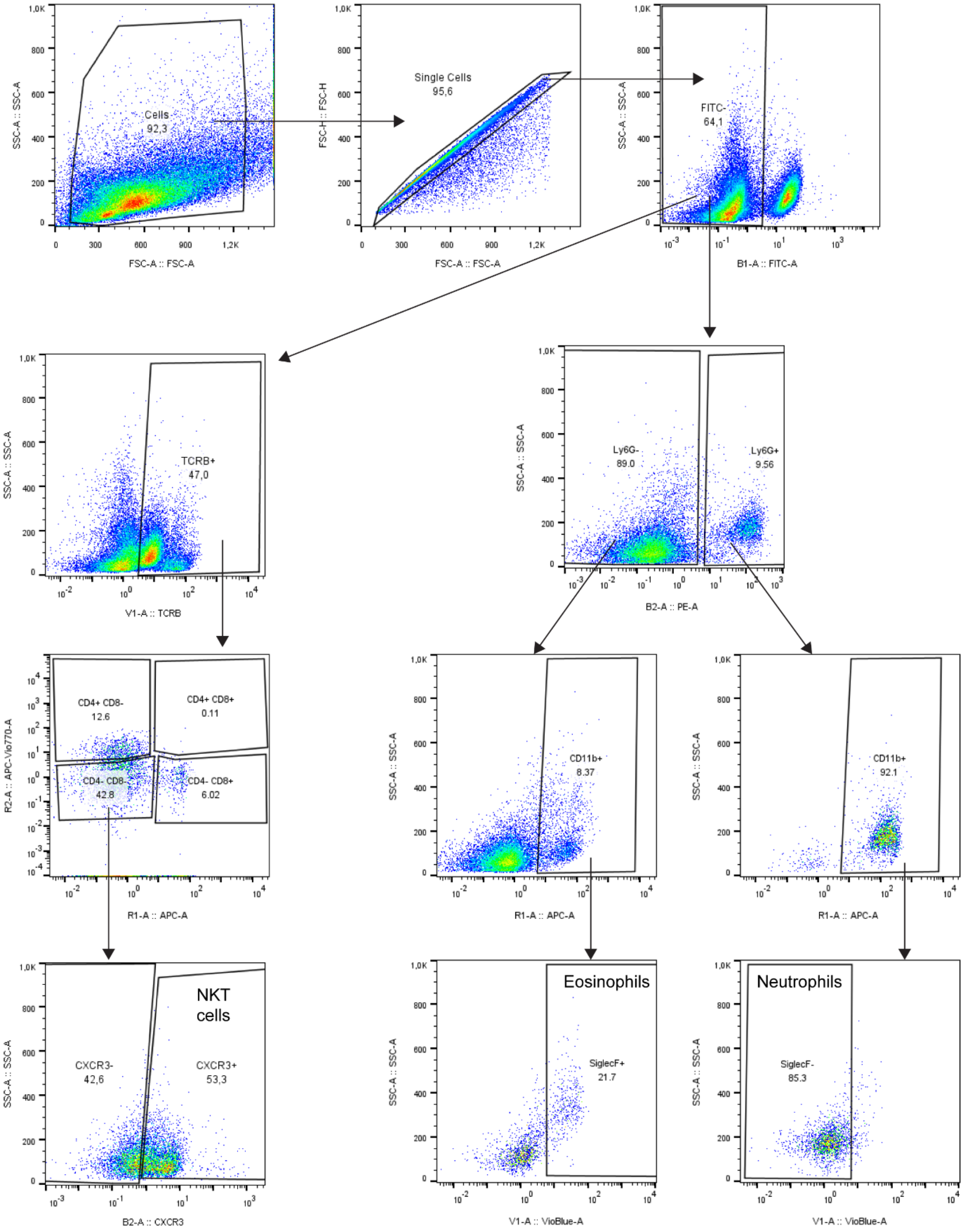


**Supplementary Figure 1. Gating strategy for NKT cells, neutrophils and eosinophils in the spleen cells.** Following singlet gating, FITC-positive tumor cells were excluded, and immune cell populations were analyzed within the GFP-negative spleen cell fraction.

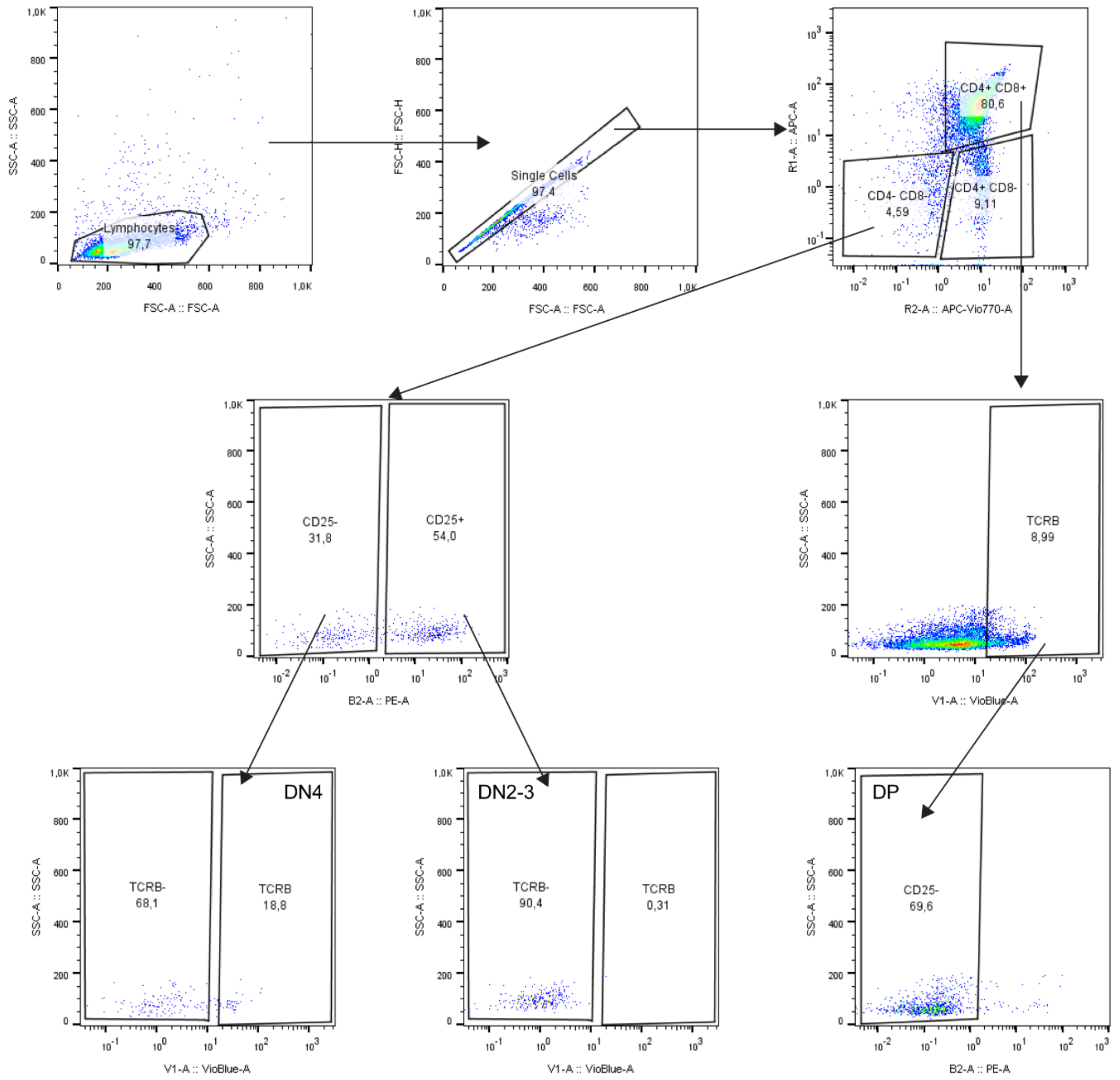


**Supplementary Figure 2. Gating strategy for DN and DP thymocytes.**

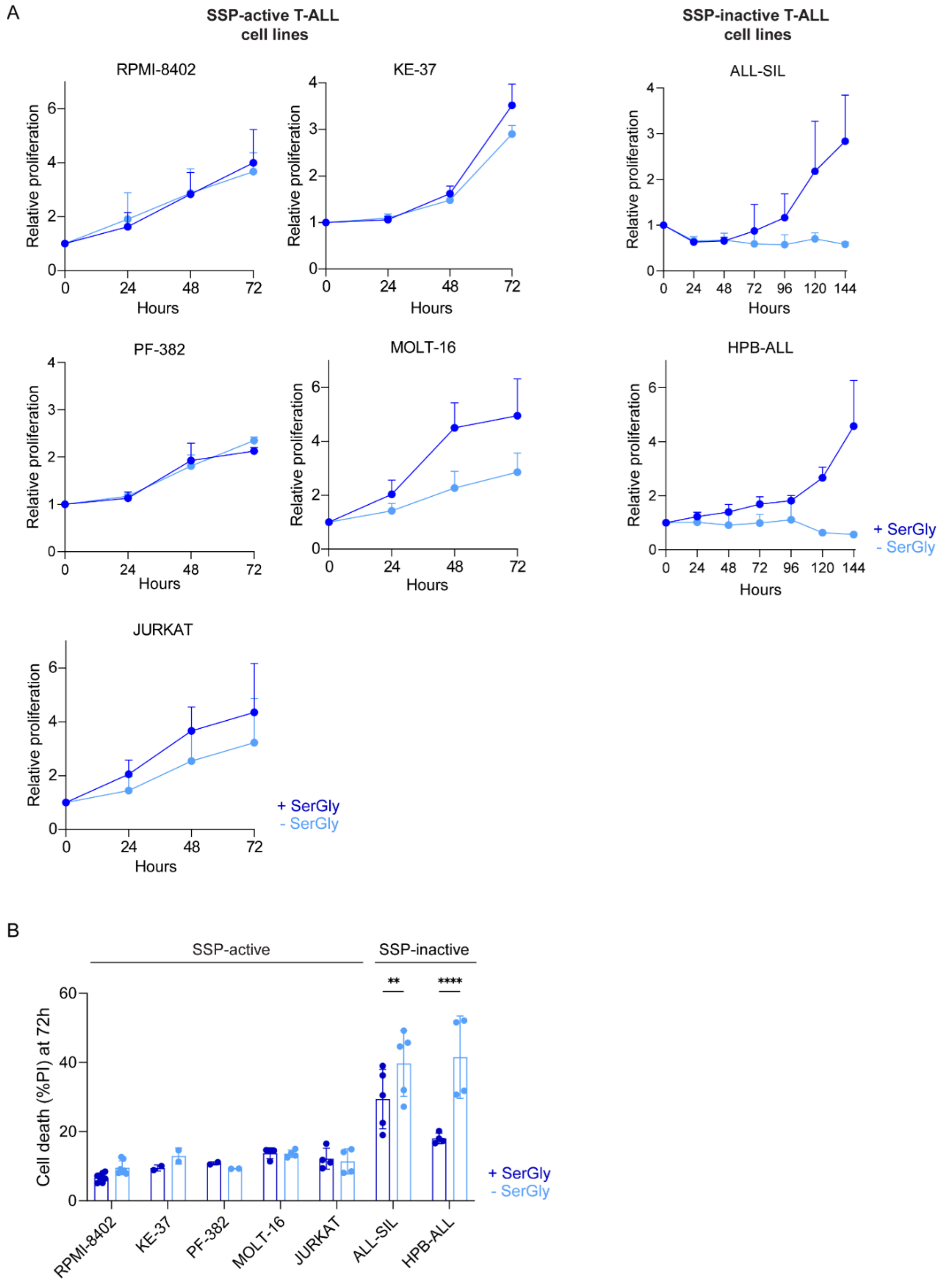


**Supplementary Figure 3. Determination of SSP-activity status of T-ALL cell lines A)** Relative cell proliferation of T-ALL cell lines RPMI-8402, KE-37, PF-382, MOLT-16, JURKAT, ALL-SIL and HPB-ALL cultured in RPMI-1640 media with or without serine and glycine, measured by flow cytometry-based cell counts over time (n≥2). **B)** Propidium iodide (PI) flow cytometry evaluating cell death induction in T-ALL cell lines RPMI-8402, KE-37, PF-382, MOLT-16, JURKAT, ALL-SIL, and HPB-ALL following 72 h of culture in RPMI-1640 medium with or without serine and glycine supplementation. The two media conditions were compared for each cell line using Two-Way ANOVA with Šidák's test for multiple comparison (n≥2). Individual dots represent independent observations. **P value < 0.01, ****P value < 0.0001.

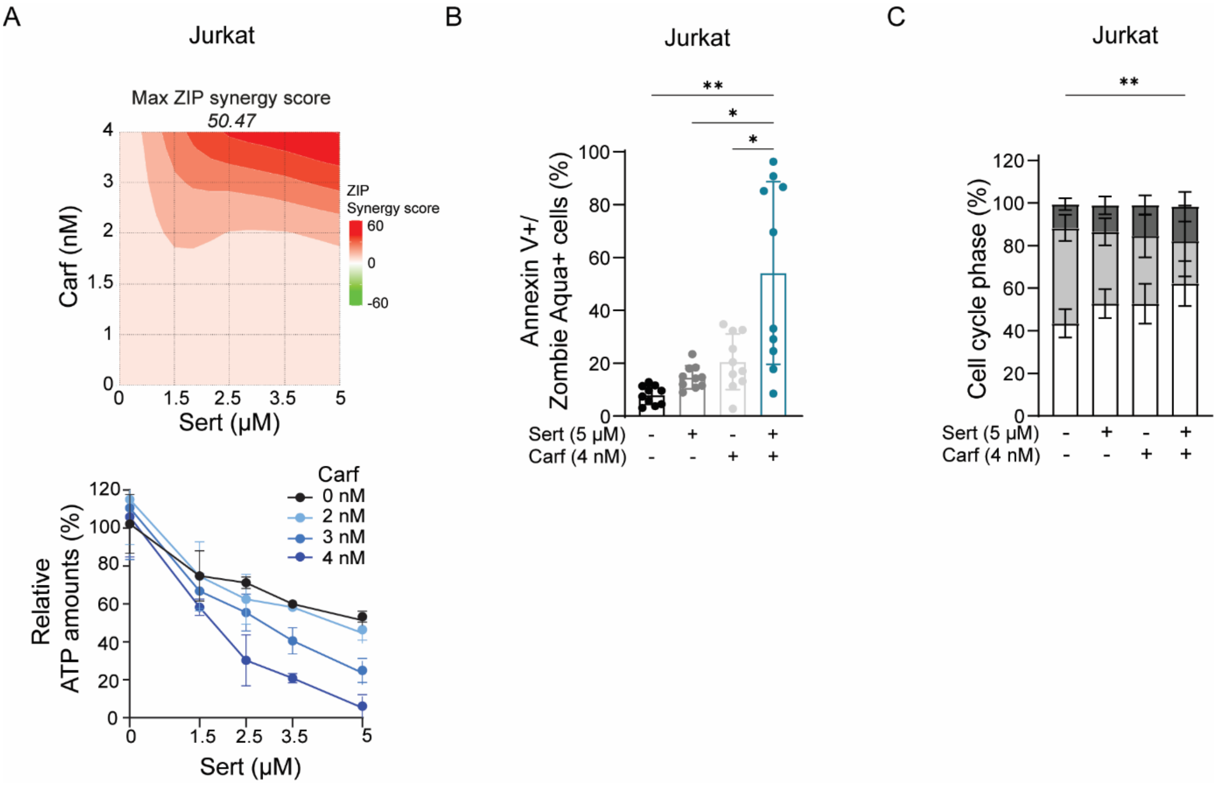


**Supplementary Figure 4. Sertraline acts synergistically with carfilzomib in SSP-active Jurkat cells. A)** Relative ATP levels measured by ATPlite assay and corresponding ZIP-score synergy map of Jurkat cells treated with sertraline, carfilzomib and their combination therapy for 48h (n≥3). A ZIP synergy score above 10 indicates synergy. **B)** Quantification of cell death by Annexin V/Zombie Aqua flow cytometry staining of Jurkat cells treated with sertraline and/or carfilzomib for 48h. The combination therapy was compared with either DMSO or single agent treated conditions using a One-Way ANOVA with Šidák's test for multiple comparison (n≥8). **C)** Quantification of BrdU-PI flow cytometry staining to analyze cell cycle distribution in Jurkat cells treated with sertraline and/or carfilzomib for 48h. S-phase contribution was compared between the different groups using Two-Way ANOVA with Turkey's test for multiple comparison (n≥6). Data are represented as mean ± standard deviation. Individual dots represent independent observations. Statistical analysis *P value < 0.05, **P value < 0.01, ***P value < 0.001, ****P value < 0.0001.

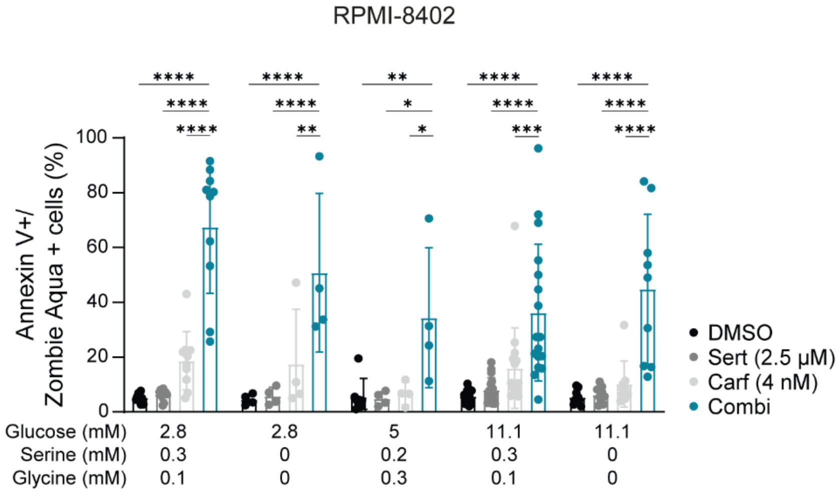


**Supplementary Figure 5. Sertraline acts synergisticlly with proteasome inhibitor carfilzomib in different culture media.** Quantification of cell death by Annexin V/Zombie Aqua flow cytometry staining in RPMI-8402 cells treated with sertraline and/or carfilzomib for 48h in RPMI-1640 media with different concentrations of glucose, serine and glycine. For each culture condition, the combination therapy was compared with either DMSO or single agent treated conditions using Two-Way ANOVA with Dunnett's test for multiple comparison (n≥4). Data are represented as mean ± standard deviation. Individual dots represent independent observations. Statistical analysis *P value < 0.05, **P value < 0.01, ***P value < 0.001, ****P value < 0.0001.

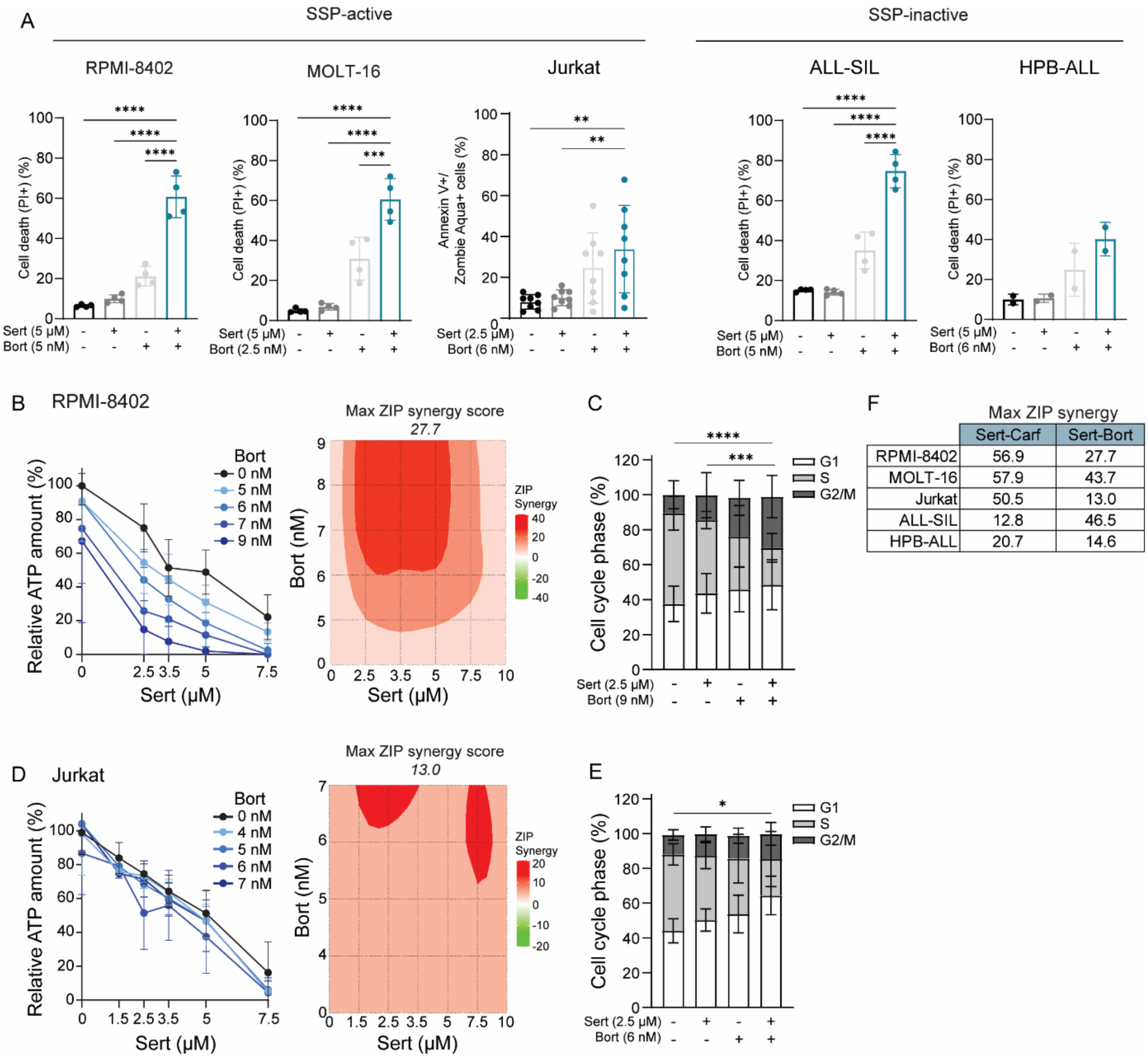


| | Sert-Carf | Sert-Bort |
|---|---|---|
| RPMI-8402 | 56.9 | 27.7 |
| MOLT-16 | 57.9 | 43.7 |
| Jurkat | 50.5 | 13.0 |
| ALL-SIL | 12.8 | 46.5 |
| HPB-ALL | 20.7 | 14.6 |

**Supplementary Figure 6. Sertraline – bortezomib combination therapy to kill T-ALL cell lines. A)** Cell death analysis measured by propidium iodide (PI) flow cytometry (RPMI-8402, MOLT-16, ALL-SIL and HPB-ALL) or Annexin V/Zombie Aqua flow cytometry (Jurkat) after 48h of treatment with DMSO, sertraline and/or bortezomib. For cell death analysis, the combination therapy condition was compared with either DMSO or single agent treated conditions using a One-Way ANOVA with Šidák's test for multiple comparison. **B)** Relative ATP amount measured by ATPlite assay (left) and corresponding ZIP score synergy map (right) of RPMI-8402 cells treated with sertraline, bortezomib and their combination therapy for 48h (n≥4). A ZIP synergy score above 10 indicates synergy. **C)** BrdU-PI cell cycle analysis of RPMI-8402 cells treated with sertraline and/or bortezomib for 48h. S-phase contribution was compared between the different treatment groups using Two-Way ANOVA with Tukey's test for multiple comparison (n=8). **D)** Relative ATP amount measured by ATPlite assay (n≥4) (left) and corresponding ZIP score synergy map (right) of Jurkat cells treated with sertraline and/or bortezomib for 48h. **E)** BrdU-PI cell cycle flow cytometry (n≥6) of Jurkat cells treated with sertraline and/or bortezomib for 48h. S-phase contribution was compared between the different groups using Two-Way ANOVA with Tukey's multiple comparison test. **F)** Overview of maximal ZIP synergy scores in RPMI-8402, MOLT-16, Jurkat, ALL-SIL and HPB-ALL for sertraline-carfilzomib versus sertraline-bortezomib combination therapy. ZIP scores for RPMI-8402 and Jurkat cells were calculated from ATPlite assay, whereas scores for the MOLT-16, ALL-SIL and HPB-ALL cell lines were calculated from PI flow cytometry data. Data are represented as mean ± standard deviation. Individual dots represent independent observations. *$P$ value < 0.05, **$P$ value < 0.01, ***$P$ value < 0.001, ****$P$ value < 0.0001.

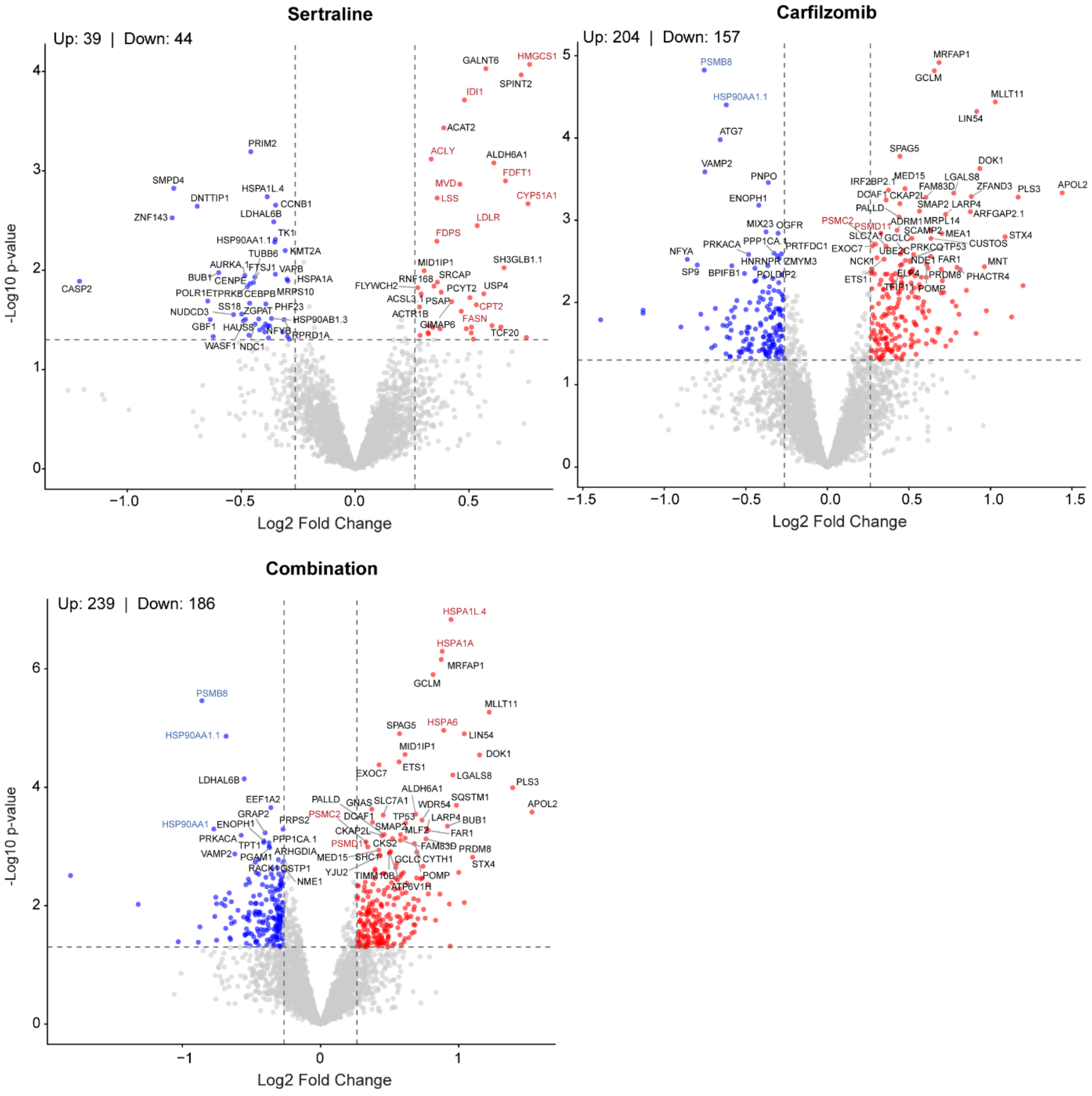


**Supplementary Figure 7. Proteomics analysis of drug treated RPMI-8402 cells.** Volcano plot of proteomics data showing significant proteins altered in RPMI-8402 cells treated with sertraline (2.5 μM), carfilzomib (4 nM) or the combination of both, each time compared to DMSO treated cells. In the sertraline graph, upregulated proteins related to cholesterol metabolism are highlighted by red protein names. In the carfilzomib and combination graphs, up- and downregulated proteins related to the proteaseome and heatshock response are highlighted by blue and red protein names.

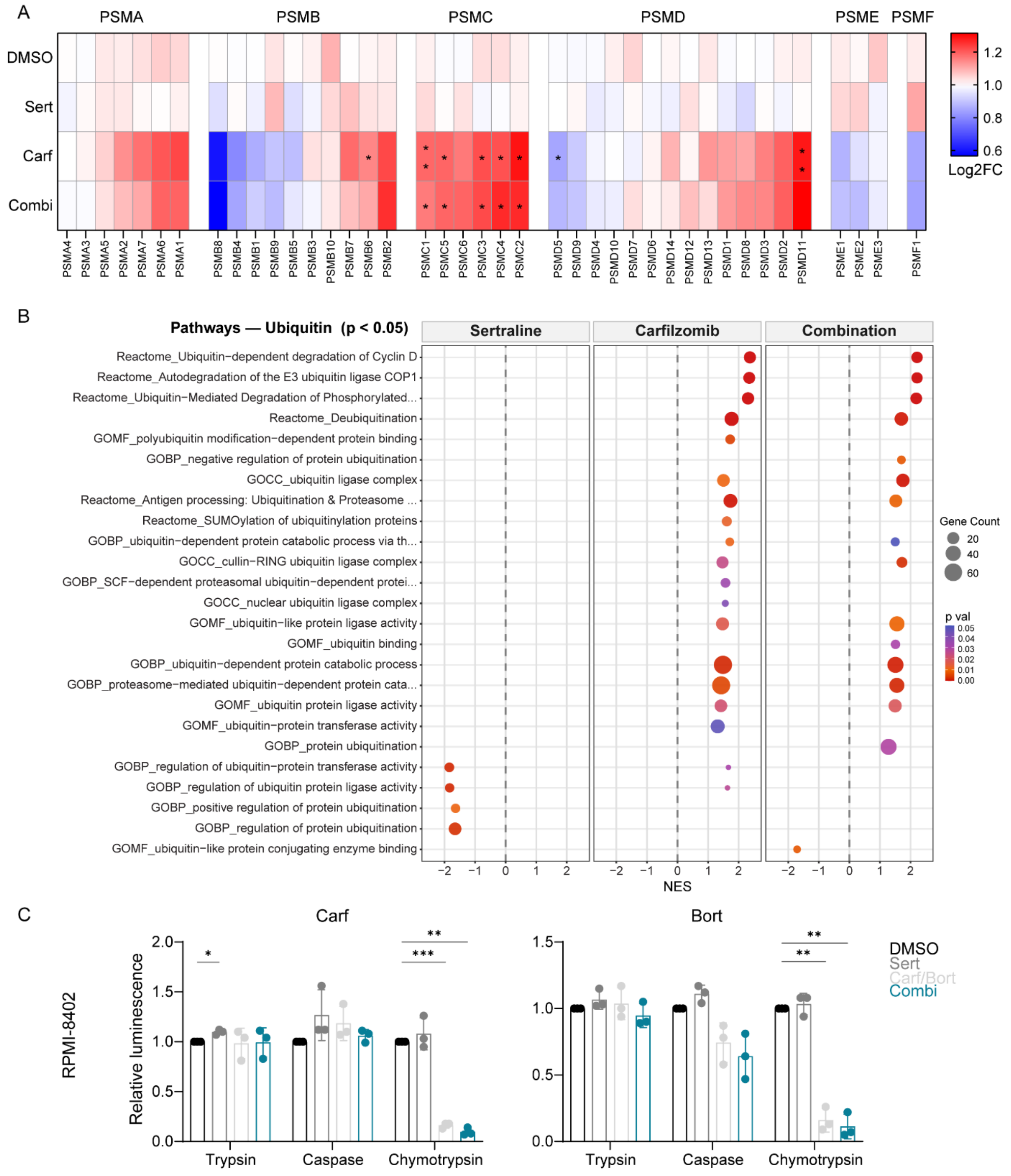


**Supplementary Figure 8. Effect of carfilzomib monotherapy on the proteasome. A)** Expression of proteasomal proteins in RPMI-8402 cells treated with DMSO, sertraline (2.5 µM) and/or carfilzomib (4 nM). The DMSO condition was compared with single agent treated or combination treated conditions using a Two-Way ANOVA with Dunnett's multiple comparison test (n=4). **B)** Gene set enrichment analysis (GSEA) bubble plot resulting from proteomics of significantly altered proteins (pval <0.05) related to ubiquitin in RPMI-8402 cells treated with sertraline and/or carfilzomib compared to DMSO for 16h. **C)** Activity of the proteasome measured by the ProteasomeGlo assay upon DMSO, sertraline, carfilzomib or bortezomib or the combination of sertraline and the proteasome inhibitors in RPMI-8402 cells. The DMSO condition was compared with single agent treated or combination treated conditions using a Two-Way ANOVA with Dunnett's multiple comparison test (n=3). Statistical analysis *P value < 0.05, **P value < 0.01, ***P value < 0.001, ****P value < 0.0001.

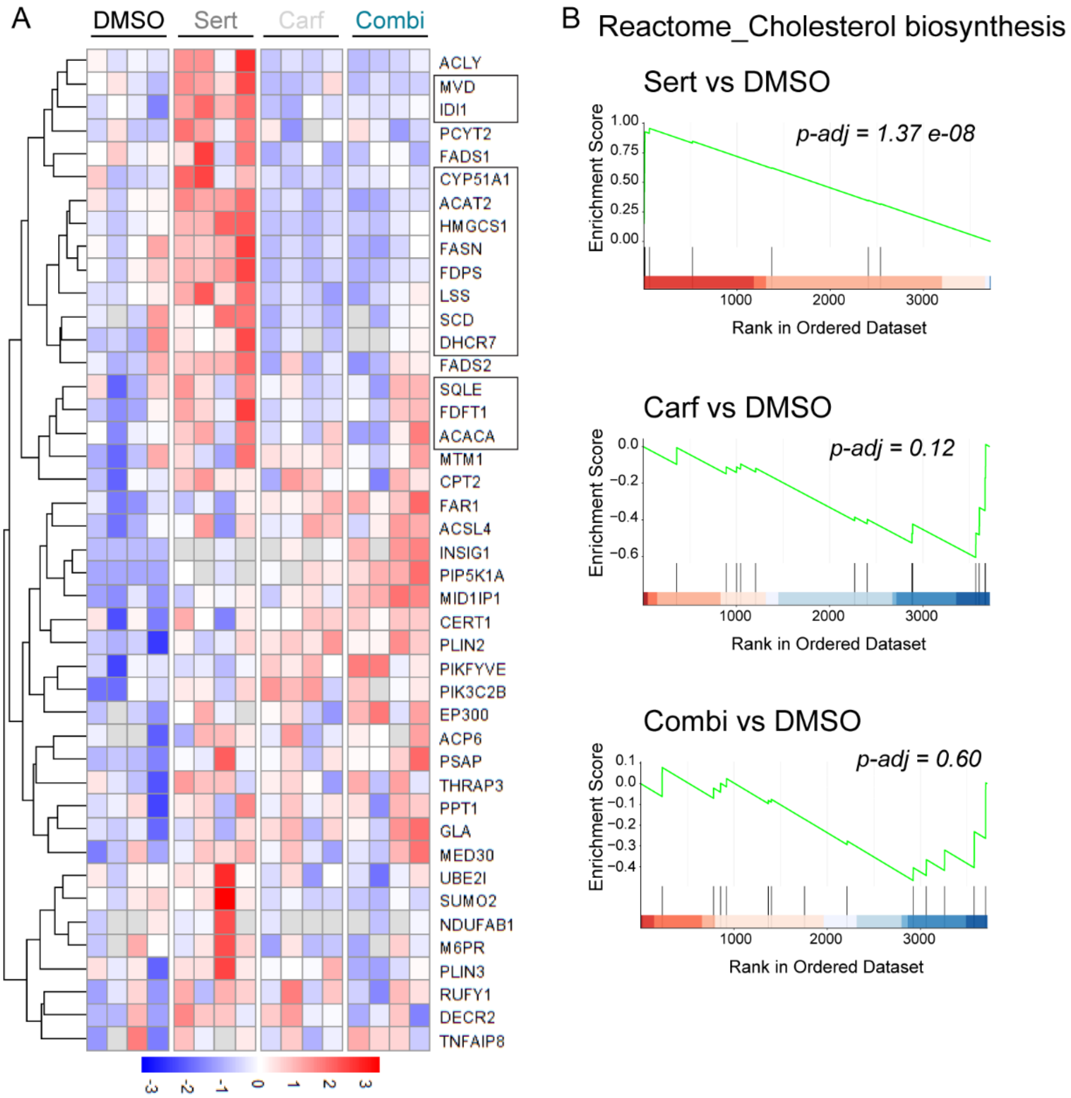


**Supplementary Figure 9. Sertraline – carfilzomib combination therapy modulates cholesterol and lipid metabolism. A)** Heat map of proteomics data showing the differentially expressed proteins related to lipid and cholesterol synthesis in RPMI-8402 cells treated with DMSO, sertraline and/or carfilzomib for 16h. Proteins indicated in the black rectangle are part of the reactome_cholesterol_biosynthesis pathway. **B)** Enrichment plot of the reactome_cholesterol_biosynthesis pathway of proteins detected in RPMI-8402 cells treated with sertraline and/or carfilzomib compared to untreated cells for 16h. NES = normalized enrichment score.

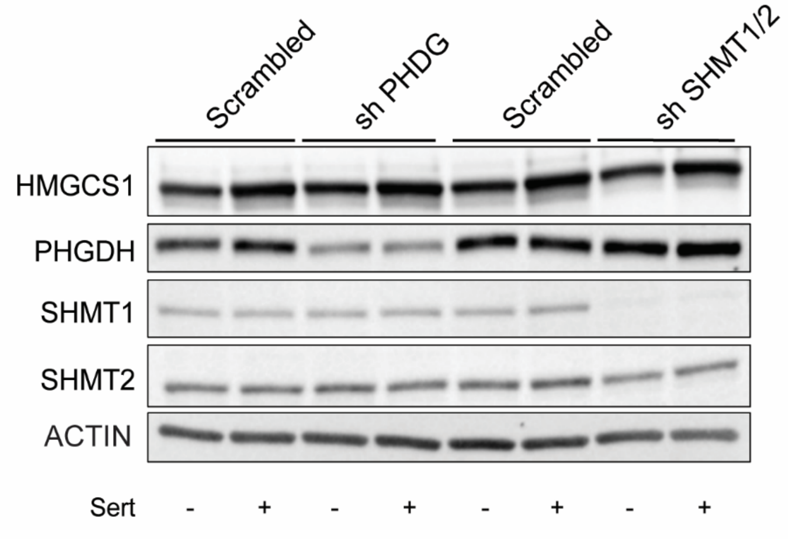


**Supplementary Figure 10. Sertraline induces cholesterol biosynthesis enzymes independently from its activity as SSP inhibitor.** Representative western blot of HMGCS1, PHGDH, SHMT1, SHMT2 and ACTIN loading control in RPMI-8402 cells transduced with scrambled shRNA, PHGDH shRNA or SHMT1/2 shRNA. Cells were treated with DMSO or 2.5 µM of sertraline for 16h. The western blot shows that sertraline treatment still leads to HMGCS1 induction, even when the SSP pathway is completely inhibited by PHGDH or SHMT1/2 knockdown.

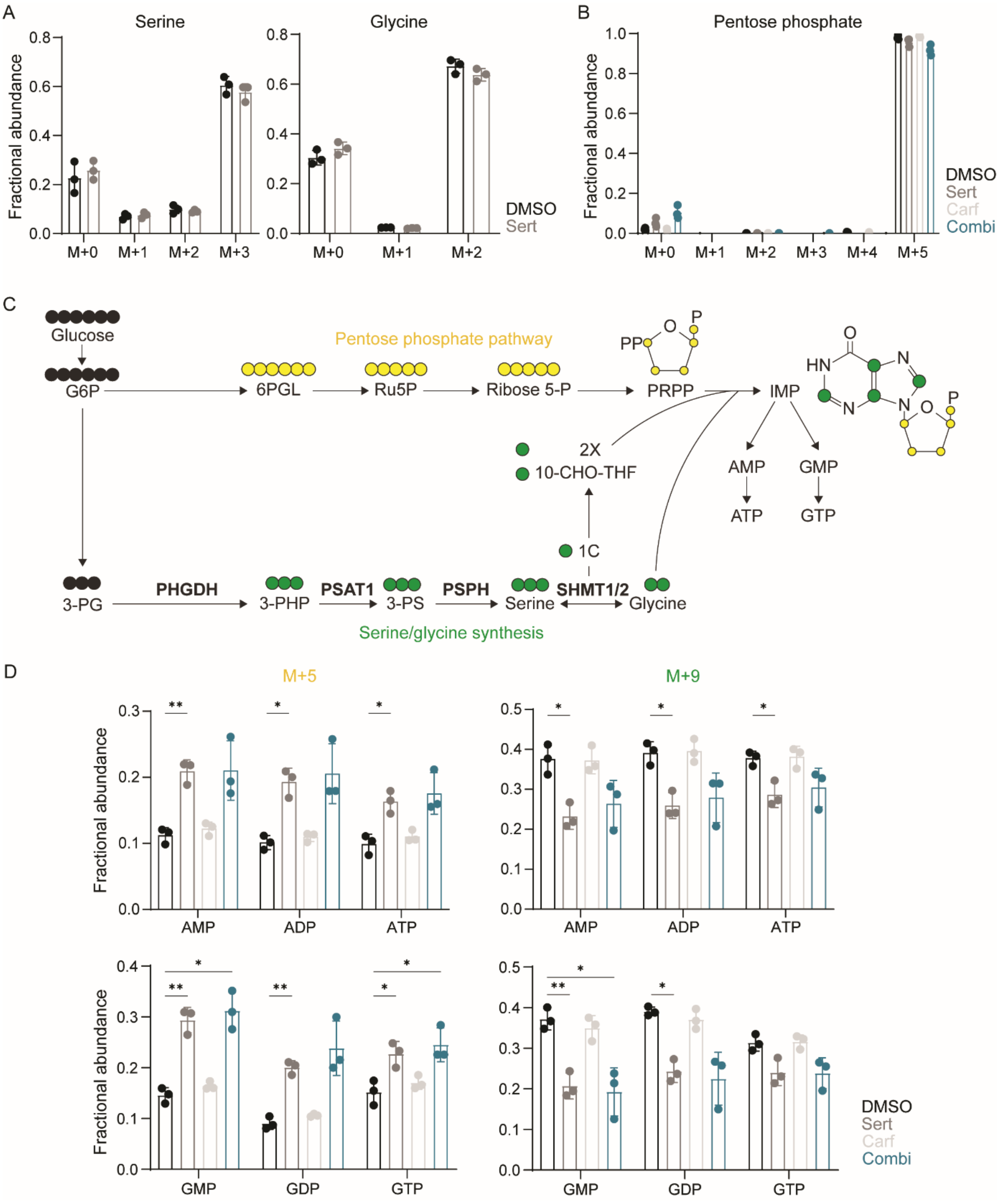


**Supplementary Figure 11. Sertraline monotherapy reduces incorporation in the SSP and downstream nucleotides. A)** Fractional abundance of 12- (M+0) and 13C-labeled (M+1-3) C-atoms in serine and glycine after treatment of RPMI-8402 cells with DMSO or sertraline (2.5 µM) for 16h. **B)** Fractional abundance of 12- (M+0) and 13C-labeled (M+1-5) C-atoms in pentose phosphate after treatment of RPMI-8402 cells with DMSO, sertraline (2.5 µM) and/or carfilzomib (4 nM) for 16h. Over 90% of pentose phosphate molecules are fully labelled (M+5). **C)** Schematic overview showing incorporation of 13C carbons from $^{13}C_6$-glucose (black) into the pentose phosphate pathway (yellow) resulting in M+5 IMP, ATP and GTP or $^{13}C_6$-glucose incorporation into the SSP (green) resulting in M+9 IMP, ATP and GTP. **D)** Fractional abundance of AMP, ADP, ATP, GMP, GDP and GTP derived from the pentose phosphate pathway (M+5) or the SSP (M+9) in RPMI-8402 cells treated with DMSO, sertraline (2.5 µM) and/or carfilzomib (4 nM) for 16h. DMSO treated condition was compared to sertraline and/or carfilzomib treated condition using Two-Way ANOVA with Dunnett's multiple comparison test (n=3). Data are represented as mean ± standard deviation. Individual dots represent independent observations. Statistical analysis *P value < 0.05, **P value < 0.01, ***P value < 0.001, ****P value < 0.0001.

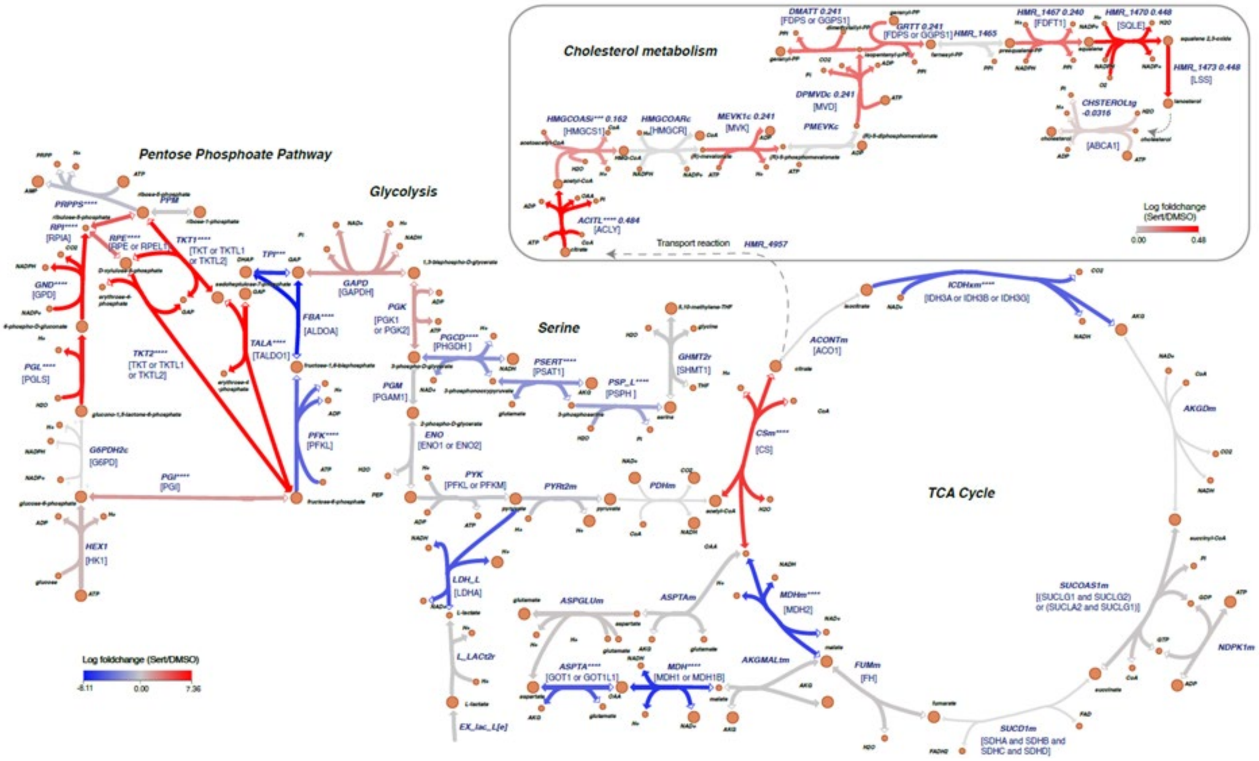


**Supplementary Figure 12. Metabolic pathway map reporting the log2-fold change of metabolic flux rates of sertraline treatment against the DMSO condition in key pathways like glycolysis, pentose phosphate pathway, TCA cycle and cholesterol metabolism pathways.** The red color represents the upregulated reactions, while the blue color represents the downregulated reactions. Statistical significance is denoted by ∗ for adjusted p < 0.05, ∗∗ for adjusted p < 0.01, ∗∗∗ for adjusted p < 0.001 and ∗∗∗∗ for adjusted p < 0.0001.

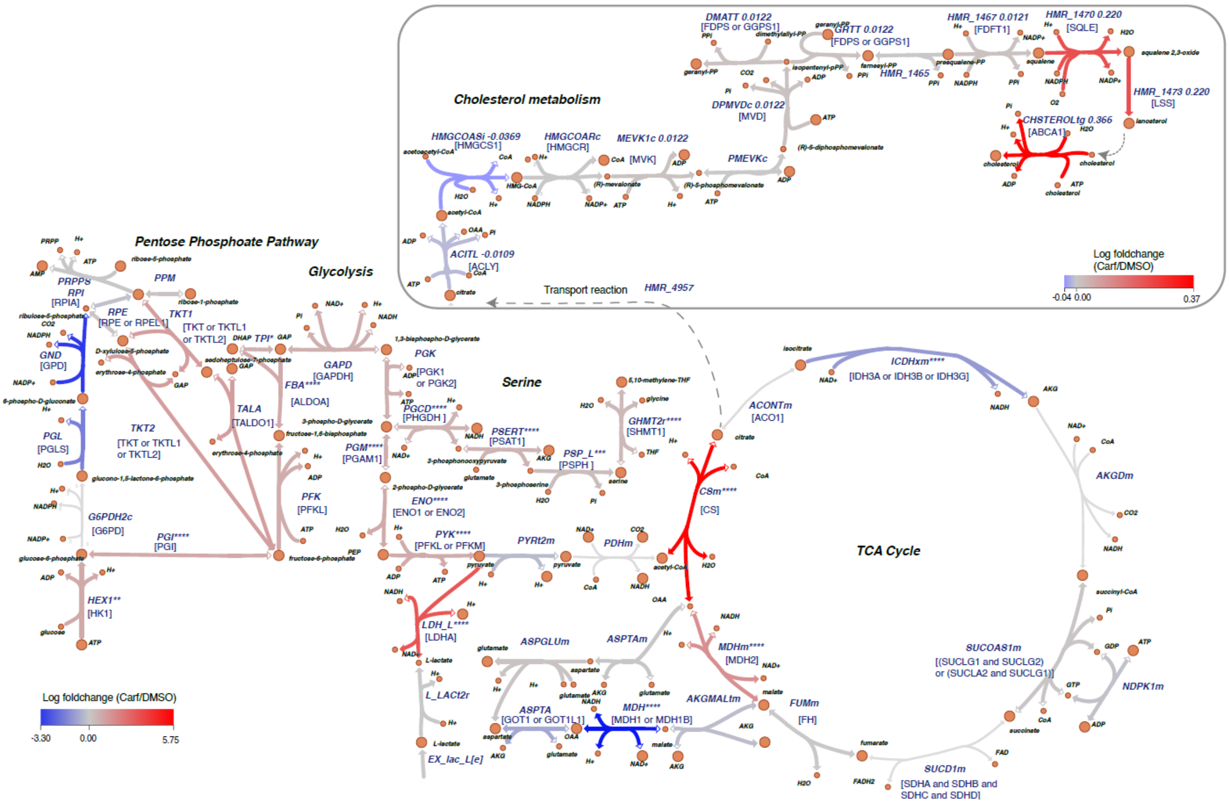


**Supplementary Figure 13. Metabolic pathway map reporting the log2-fold change of metabolic flux rates of carfilzomib treatment against the DMSO condition in key pathways like glycolysis, pentose phosphate pathway, TCA cycle and cholesterol metabolism pathways.** The red color represents the upregulated reactions, while the blue color represents the downregulated reactions. Statistical significance is denoted by $*$ for adjusted $p < 0.05$, $**$ for adjusted $p < 0.01$, $***$ for adjusted $p < 0.001$ and $****$ for adjusted $p < 0.0001$.

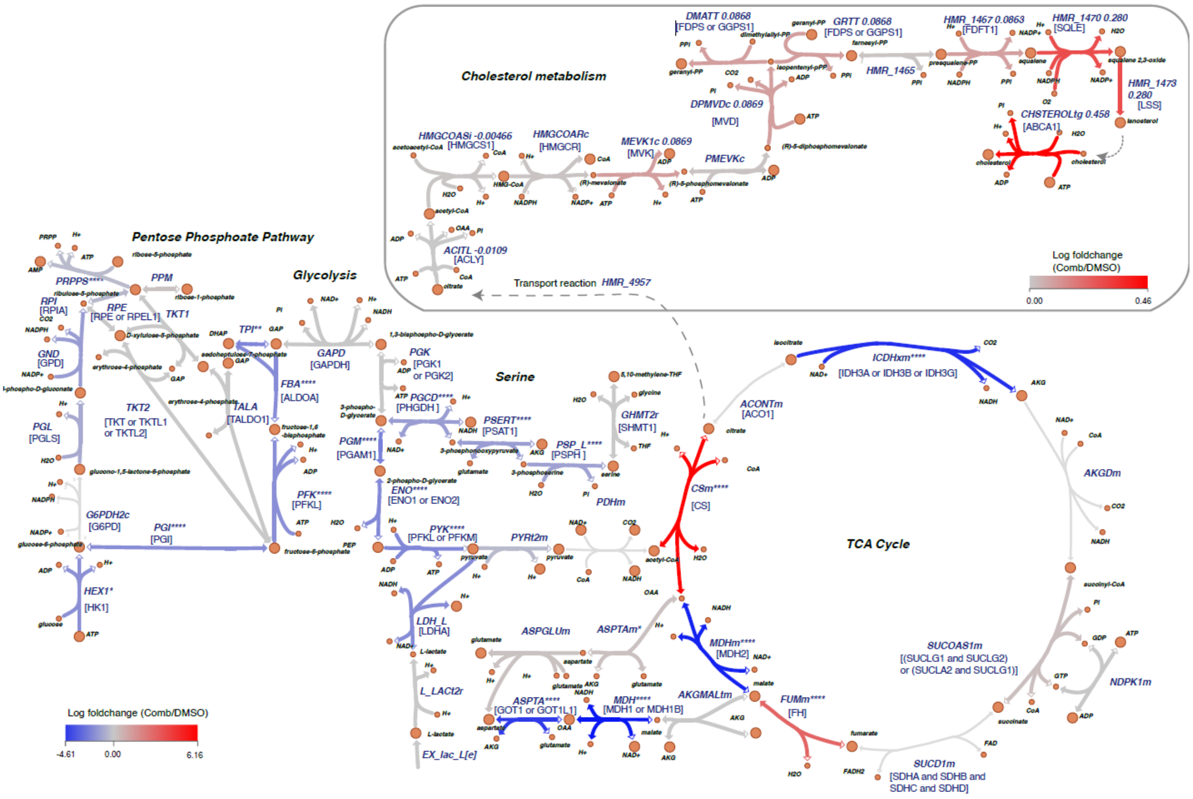


**Supplementary Figure 14. The metabolic pathway map reporting the log2-fold change of metabolic flux rates of combined treatment against the DMSO condition in key pathways like glycolysis, pentose phosphate pathway, TCA cycle and cholesterol metabolism pathways.** The red color represents the upregulated reactions, while the blue color represents the downregulated reactions. Statistical significance is denoted by ∗ for adjusted $p < 0.05$, ∗∗ for adjusted $p < 0.01$, ∗∗∗ for adjusted $p < 0.001$ and ∗∗∗∗ for adjusted $p < 0.0001$.

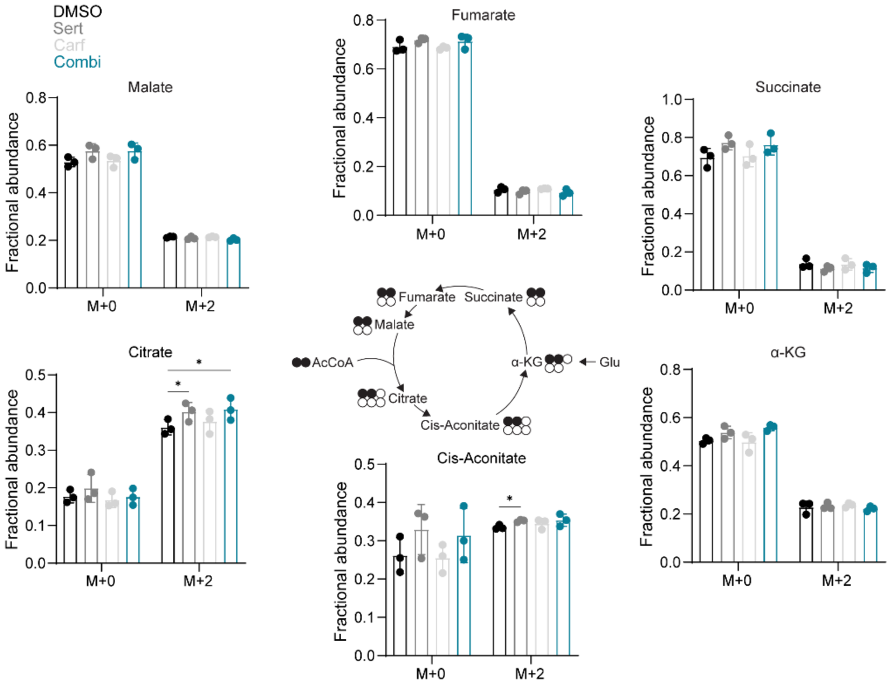


**Supplementary Figure 15. Effect of sertraline – carfilzomib combination therapy on 13C incorporation in the TCA cycle.** Fractional abundance of 12- (M+0) and 13C-labeled (M+2) C-atoms in TCA intermediates after treatment of RPMI-8402 cells with DMSO, sertraline (2.5 µM) and/or carfilzomib (4 nM) for 16h. The DMSO treated condition was compared to the sertraline and/or carfilzomib treated condition for each isotopologue using Two-Way ANOVA with Dunnett's multiple comparison test (n=3). Data are represented as mean ± standard deviation. Individual dots represent independent observations. Statistical analysis *P value < 0.05, **P value < 0.01, ***P value < 0.001, ****P value < 0.0001.

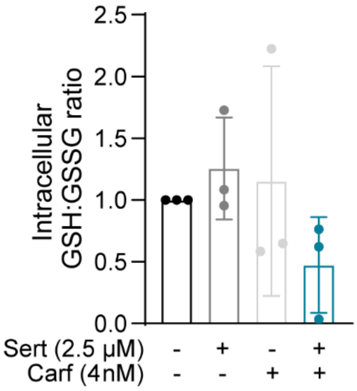


**Supplementary Figure 16. Sertraline – carfilzomib treatment induces cellular stress.** Ratio of reduced glutathione (GSH) to oxidized GSH (GSSG) in RPMI-8402 cells treated with DMSO, sertraline and/or carfilzomib for 16h. The DMSO treated condition was compared to the sertraline and/or carfilzomib treated condition using One-Way ANOVA with Dunnett's multiple comparison test (n=3). Data are represented as mean ± standard deviation. Individual dots represent independent observations. Statistical analysis *P value < 0.05, **P value < 0.01, ***P value < 0.001, ****P value < 0.0001.

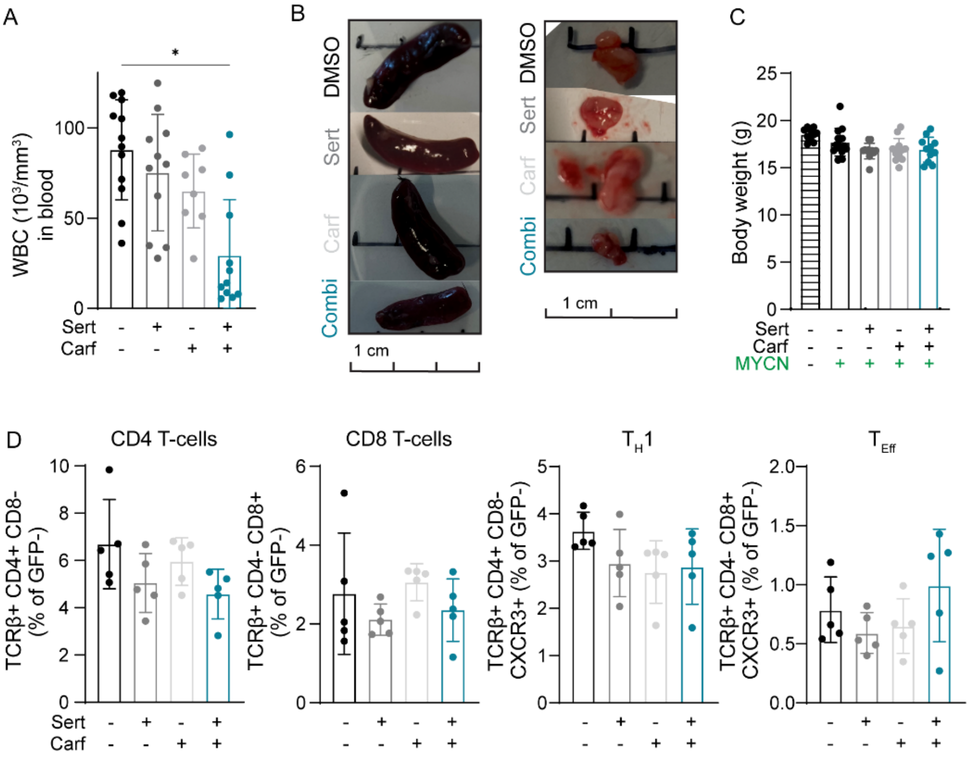


**Supplementary Figure 17. Effects of sertraline – carfilzomib combination therapy in MYCN-PTCL *in vivo* model. A)** Effect of DMSO, sertraline and/or carfilzomib therapy on the total white blood cell counts (WBC) in the blood of PTCL mice at disease end stage (day 13). **B)** Representative images of the spleen (left) and thymus (right) upon DMSO, sertraline and/or carfilzomib treatment of MYCN-PTCL mice. **C)** Effect of the mono- and combination therapies on body weight compared to healthy mice. **D)** Effect of sertraline – carfilzomib combination therapy on percentage of CD4 single positive T-cells, single positive CD8 T-cells, T-helper 1 cells and T-effector cells in spleen analyzed by flow cytometry in the GFP-negative population. Data are represented as mean ± standard deviation. Individual dots represent independent observations. Statistical analysis was done using a One-Way ANOVA with Dunnett's (A,D) or Tamhane T2 (C) multiple comparison test (n≥3) whereby the DMSO condition was compared to others. Statistical analysis *P value < 0.05, **P value < 0.01, ***P value < 0.001, ****P value < 0.0001.

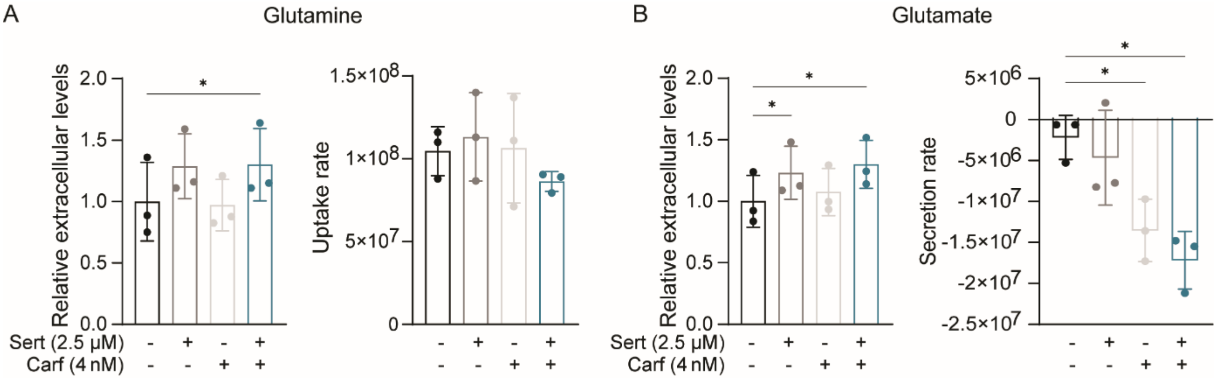


**Supplementary Figure 18. Extracellular glutamine and glutamate levels in T-ALL cells treated with sertraline – carfilzomib therapy. A-B)** Relative extracellular levels of glutamine (A) and glutamate (B) and their uptake or secretion rate in RPMI-8402 cells treated with sertraline and/or carfilzomib for 16h. The DMSO treated condition was compared to the sertraline and/or carfilzomib treated condition using One-Way ANOVA with Dunnett's multiple comparison test (n=3). Data are represented as mean ± standard deviation. Individual dots represent independent observations. Statistical analysis *P value < 0.05, **P value < 0.01, ***P value < 0.001, ****P value < 0.0001.

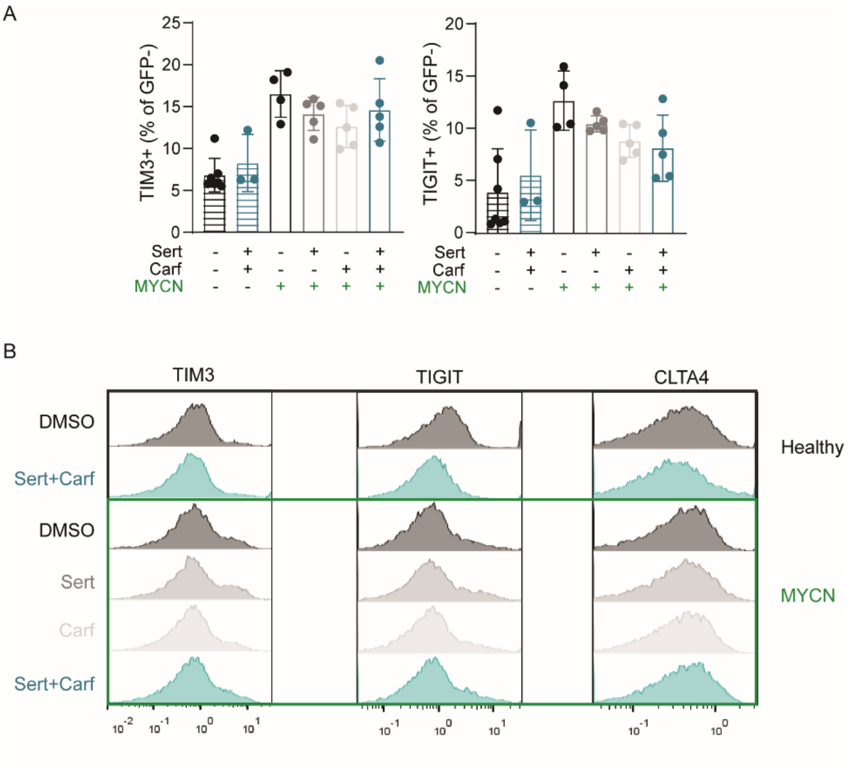


**Supplementary Figure 19. Effects of sertraline – carfilzomib combination therapy in MYCN-PTCL *in vivo* model. A-B)** TIM3, TIGIT and CTLA-4 expression in the GFP-negative population of spleen cells of healthy or T-cell lymphoma mice treated with vehicle, sertraline, carfilzomib or the combination. Representative histograms are shown in (B). Data are represented as mean ± standard deviation. Individual dots represent independent observations. Statistical analysis was done using a One-Way ANOVA with Dunnett's multiple comparison test (n≥3) whereby the DMSO condition was compared to others. Statistical analysis *P value < 0.05, **P value < 0.01, ***P value < 0.001, ****P value < 0.0001.

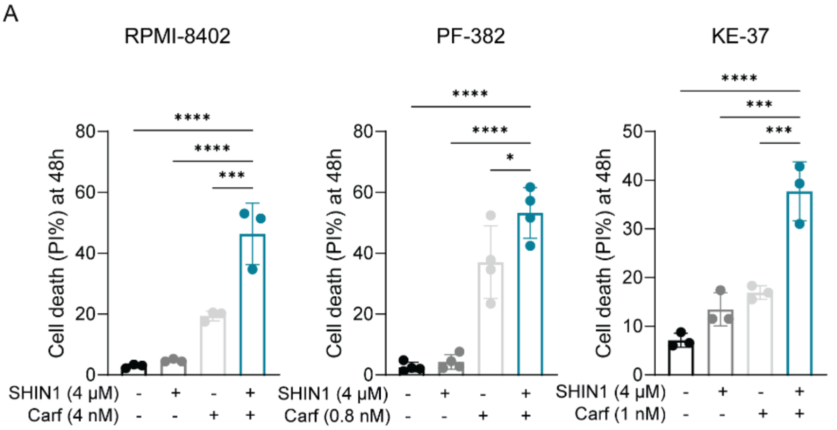


B

Max ZIP synergy

| | Sert-Carf | SHIN1-Carf |
|---|---|---|
| RPMI-8204 | 44.5 | 30.6 |
| PF-382 | 42.7 | 19.2 |
| KE-37 | 64.4 | 17.7 |

**Supplementary Figure 20. Synergy between sertraline – carfilzomib versus SHIN1 – carfilzomib in SSP-active T-ALL cell lines.** Cell death analysis measured by PI flow cytometry staining of SSP-active RPMI-8402, PF-382 and KE-37 T-ALL cell lines after 48h of treatment with DMSO, SHMT inhibitor SHIN1 and/or carfilzomib. The combination therapy was compared with either DMSO or single agent treated conditions using a One-Way ANOVA with Šidák's test for multiple comparison. Data are represented as mean ± standard deviation. Individual dots represent independent observations. *P value < 0.05, **P value < 0.01, ***P value < 0.001, ****P value < 0.0001. **B)** Overview of maximal ZIP synergy scores in RPMI-8402, PF-382 and KE-37 for sertraline-carfilzomib versus SHIN1-carfilzomib combination therapy.

| CXCL8 | | | |
|---|---|---|---|
| DMSO | Sert | Carf | Combi |
| NA | NA | NA | 65338.5 |
| NA | NA | NA | 88072.5 |
| NA | NA | NA | 74969 |
| NA | NA | NA | 235521 |

**Supplementary Figure 21. CXCL8 detection by label-Free Quantification (LFQ) mass-spectrometry**. LFQ intensity values for DMSO, Sertraline (Sert), carfilzomib (Carf) and combination RPMI-8402 treated cells.

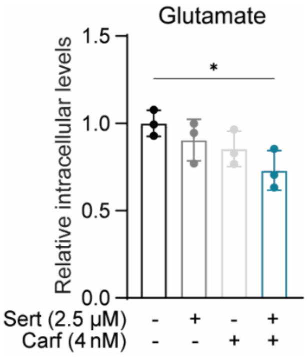


**Supplementary Figure 22. Effects of sertraline – carfilzomib combination therapy on intracellular glutamate levels.** Relative intracellular glutamate levels in RPMI-8402 cells treated with DMSO, sertraline and/or carfilzomib for 16h. The DMSO treated condition was compared to sertraline and/or carfilzomib treated condition using One-Way ANOVA (n≥3). *P value < 0.05

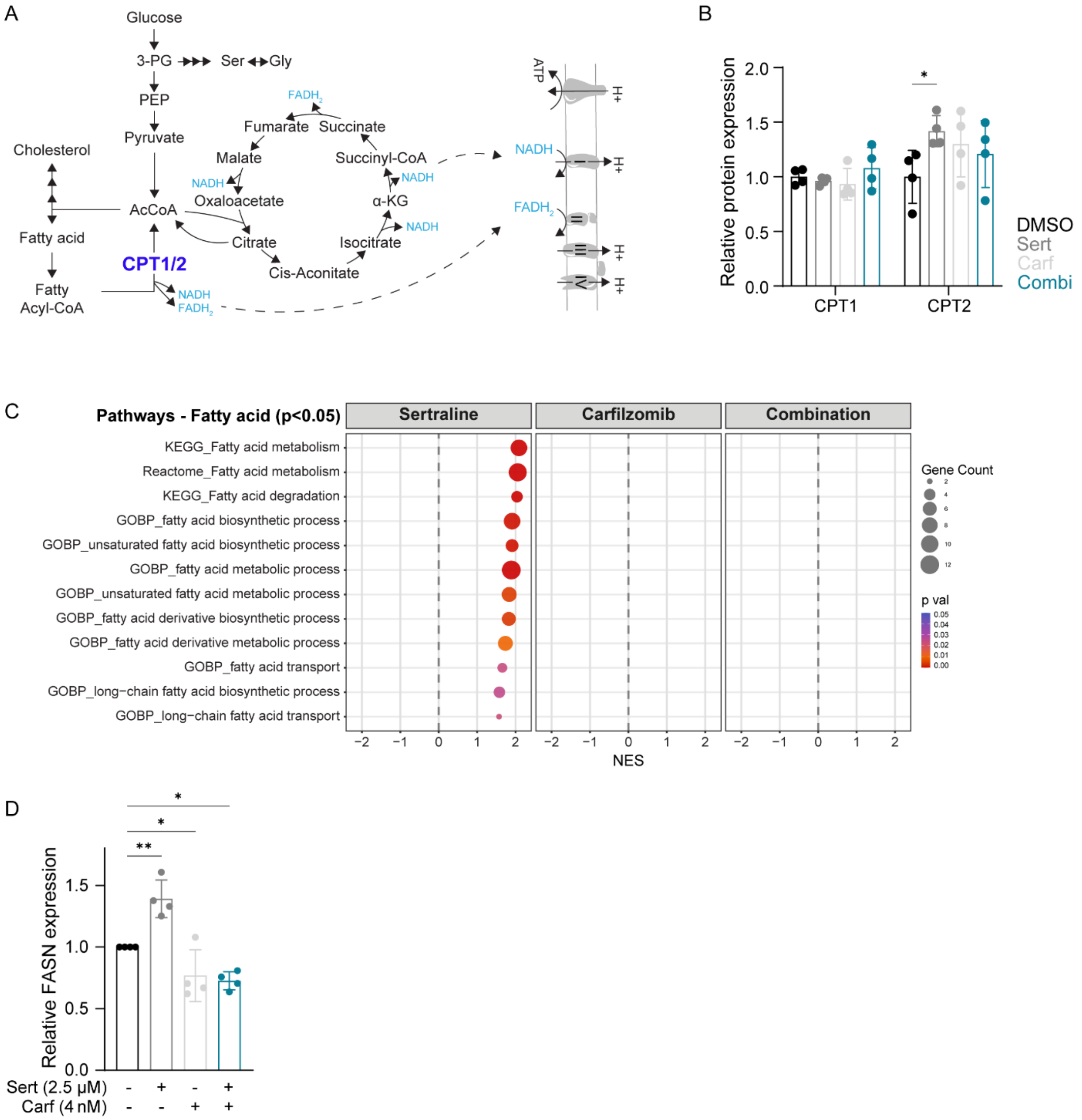


**Supplementary Figure 23. Sertraline monotherapy alters fatty acid metabolism and oxidation. A)** Scheme of NADH and $FADH_2$ donation to the electron transport chain coming from the TCA cycle or fatty acid oxidation catalyzed by CPT1/CPT2. **B)** Relative protein expression of CPT1 and CPT2 in RPMI-8402 cells treated with sertraline and/or carfilzomib therapy for 16h derived from proteomics data. For each protein, the DMSO condition was compared to either single agent or combination treatment using Two-Way ANOVA with Dunnett's multiple comparison test (n=4). **C)** Gene set enrichment analysis (GSEA) bubble plot resulting from proteomics of significantly altered proteins (pval <0.05) related to fatty acids in RPMI-8402 cells treated with sertraline and/or carfilzomib compared to DMSO for 16h. **D)** Relative intracellular expression of FASN in RPMI-8402 cells treated with sertraline and/or carfilzomib for 16h derived from proteomics data. The DMSO condition was compared to single agent or combination treatment using One-Way ANOVA with Dunnett's multiple comparison test (n=4). Data are represented as mean ± standard deviation. Individual dots represent independent observations. Statistical analysis *P value < 0.05, **P value < 0.01, ***P value < 0.001, ****P value < 0.0001.

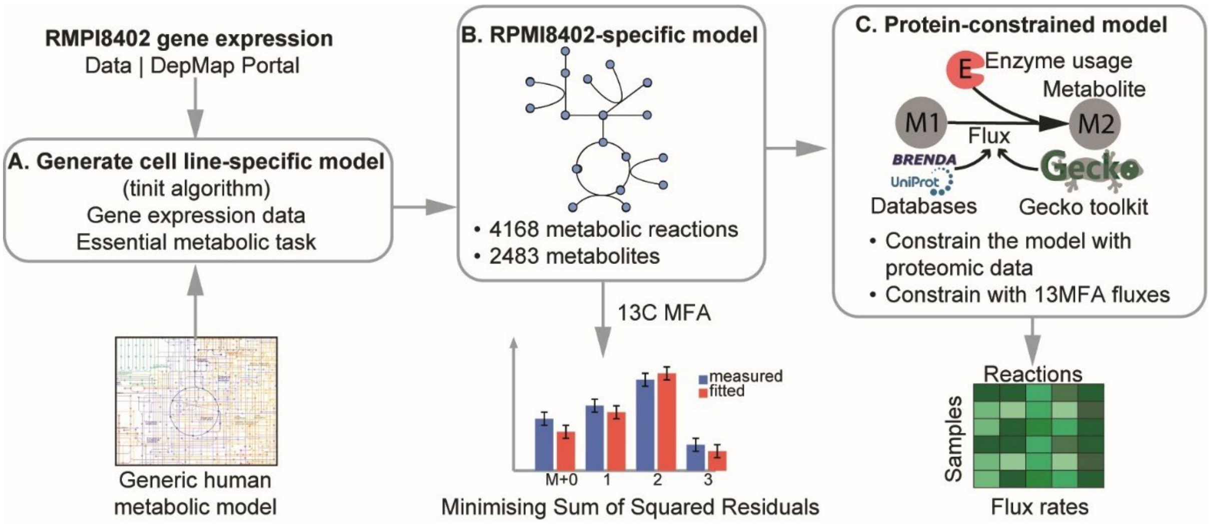


**Supplementary Figure 24. Workflow for estimating condition-specific flux rates, integrating 13C-glucose tracing isotopomer and proteomics data**. **A)** The RPMI-8402 cell-line-specific metabolic model was constructed leveraging cell-line gene expression data from the DepMap portal and the Human-Gem metabolic model. **B)** The 13C glucose carbon tracing data were used to estimate the 13C MFA for reactions in central carbon metabolism. **C)** The GECKO toolkit was applied to use the 13C MFA flux estimates for constraining the reactions in the whole RPMI-8402 cell-line-specific metabolic model, along with proteomics data.

### SUPPLEMENTARY TABLES

#### Supplementary Table 1. List of cell lines.

| Cell line | Source | Disease subtype |
|---|---|---|
| KE-37 | DSMZ | T-ALL |
| MOLT-16 | DSMZ | T-ALL |
| ALL-SIL | DSMZ | T-ALL |
| HPB-ALL | DSMZ | T-ALL |
| Jurkat | ATCC | T-ALL |
| RPMI-8402 | DMSZ | T-ALL |
| PF-382 | DMSZ | T-ALL |

#### Supplementary Table 2. Compounds.

| Compound | Company | Cat# | Dissolved in |
|---|---|---|---|
| Bortezomib | TebuBio | 10-2120 | DMSO |
| Carfilzomib | MedChemExpress | HY-10455 | DMSO |
| Lovastatin | MedChemExpress | HY-N0504 | DMSO |
| Sertraline | Sigma-Aldrich | S6319 | DMSO |
| SHIN1 | MedChemExpress | HY-112066 | DMSO |
| NCT503 | MedChemExpress | HY-101966 | DMSO |

#### Supplementary Table 3. Antibodies used for flow cytometry.

| Target | Fluorophore | Company | Cat# |
|---|---|---|---|
| Annexin V | PE | Biolegend | 640947 |
| Cleaved caspase 3 | / | Cell Signaling Technology | 9664S |
| Rabbit IgG (H+L) | Alexa Fluor 488 | Invitrogen | A11070 |

| | | | |
|---|---|---|---|
| BrdU | FITC | Biolegend | 364104 |
| TCR-β | Brilliant Violet 421 | Biolegend | 109230 |
| CD4 | APC/Fire 750 | Biolegend | 100460 |
| CD8a | APC | Biolegend | 100712 |
| CD25 | PE | Biolegend | 102007 |
| CXCR3 | PE | Biolegend | 126506 |
| Ly6G | PE | Biolegend | 164503 |
| CD11b | APC | Biolegend | 101212 |
| SiglecF | Brilliant Violet 421 | Biolegend | 155509 |
| Ly6C | APC/Fire 750 | Biolegend | 128046 |
| CD80 | Brilliant Violet 421 | Biolegend | 104726 |
| NK1.1 | APC/Fire 750 | Biolegend | 108752 |
| CD206 | PE | Biolegend | 141706 |
| CD11c | PE | Biolegend | 117307 |
| MHCII | APC/Fire 750 | Biolegend | 107652 |
| XCR1 | Brilliant Violet 421 | Biolegend | 148216 |
| CD45R | Brilliant Violet 421 | Biolegend | 103251 |
| CD19 | APC/Fire 750 | Biolegend | 115558 |
| TIM-1 | PE | Miltenyi | 130-110-325 |
| CD138 | APC | Miltenyi | 130-122-945 |
| PD-1 | PE | Biolegend | 109103 |
| CTLA-4 | APC | Miltenyi | 130-116-455 |
| TIM-3 | APC/Fire 750 | Biolegend | 119738 |
| TIGIT | Brilliant Violet 421 | Biolegend | 142111 |
| y-H2AX (Ser139) | / | Cell Signaling Technology | 9718 |

**Supplementary Table 4. Dyes used for flow cytometry.**

| Dye | Company | Cat# |
|---|---|---|
| Zombie Aqua | Biolegend | 423102 |
| Propidium iodide | Sigma-Aldrich | 81845 |
| BrdU | eBioscience | 00-4440-51A |
| CellROX Deep Red | ThermoFisher Scientific | C10422 |
| Nile Red | Cayman chemical | 500001 |

**Supplementary Table 5. Antibodies used for Western blot.**

| Target | Company | Cat# |
|---|---|---|
| y-H2AX (Ser139) | Cell Signaling Technology | 9718 |
| NKX2-1 | Abcam | ab133737 |
| HMGCS1 | Cell Signaling Technology | 42201S |
| SQLE | Cell Signaling Technology | 40659S |
| ACLY | Cell Signaling Technology | 4332S & 13390 |
| PHGDH | Cell Signaling Technology | 66350 |
| PSPH | ProteinTech | 14513-1-AP |
| PSAT1 | ProteinTech | 10501-1-AP |
| SHMT1 | Abcam | ab186130 |
| SHMT2 | ProteinTech | 11099-1-AP |
| VINCULIN | Sigma-Aldrich | V9131 |
| β-ACTIN | Sigma-Aldrich | A1978 |
| Anti-Rabbit IgG-HRP | ThermoFisher Scientific | 31462 |
| Anti-Mouse IgG-HRP | ThermoFisher Scientific | 31432 |

**Supplementary Table 6. shRNA sequences.**

| Target | shRNA sequence | Plasmid backbone |
|---|---|---|
| Scrambled control | CCCTTCTCCGGAACGTGTCACGTTTCAAGAGAACGTGGACACGGTTCGGAGAATTTA | pLKO1 |
| PHGDH | CCGGGCTTCGATGAAGGACGGCAAACTCGAGTTTGCCGTCCTTCATCGAAGCTTTTT | pLKO1 |
| Scrambled control | CCTAAGGTTAAGTCGCCCTCGCTCGAGCGAGGGCGACTTAACCTTAGG | pLV |
| SHMT1 | CCTAGGCTCTTGCTTAAATAACTCGAGTTATTTAAGCAAGAGCCTAGG | pLV |
| SHMT2 | CGGAGAGTTGTGGACTTTATACTCGAGTATAAAGTCCACAACTCTCCG | pLV |

**Supplementary Table 7 (See separate Excel file). Metabolomics analysis of the serum of PTCL and healthy mice** (accompanying data of Figure 8G).

**Supplementary Table 8 (See separate Excel file). Accompanying data of metabolic RPMI-8402 AI model.**

### SUPPLEMENTARY METHODS

#### 1. Generation of metabolic RPMI-8402 AI model

To investigate metabolic alterations in T-cell acute lymphoblastic leukemia (T-ALL) across treatment conditions, we estimated metabolic reaction flux rates using a three-step approach that integrates 13C-glucose isotopomer data with protein expression levels (**Supplementary Figure 24**). First, using the FreeFlux framework[1], we estimated steady-state 13C-based fluxes for the reactions in central carbon metabolism based on 13C-glucose labelling patterns. Then, these flux estimates were used to constrain the reaction bounds of the RPMI-8402 cell line-specific metabolic model (manually curated and reconstructed from the Human-GEM model, Robinson et al., 2020). Finally, proteomics data were integrated into the constrained RPMI-8402 model to infer flux distributions under each treatment condition[2].

#### 1.1. Construction of RPMI-8402 cell line-specific genome-scale metabolic model

To map the metabolic alterations in T-cell acute lymphoblastic leukemia (T-ALL), we first constructed the RPMI-8402 cell-line-specific Genome-Scale Metabolic Model (GSMM) by leveraging the highly curated Human-GEM model[3], which provides comprehensive mechanistic insight into cellular metabolism, enabling detailed comparisons between treated and control groups.

Starting from the general Human-GEM model, we removed the reactions that were not directly relevant to RPMI-8402 cell line metabolism or reflected T-ALL metabolism, and obtained an RPMI-8402 cell line-specific model. This step was performed using the task-driven Integrative Network Inference for Tissues (tINIT) algorithm, which is specifically designed to construct context-specific metabolic models by integrating RPMI-8402 cell line-specific gene expression data, derived from the Depmap portal[4], with a generic genome-scale metabolic network. First, gene expression profiles were mapped to the reactions in the Human-GEM model through established gene-protein-reaction (GPR) associations. Each reaction was assigned a confidence score based on the expression level of its associated genes. In parallel, a set of essential metabolic tasks relevant to core cellular functions was defined to ensure that the reconstructed models remained biologically feasible. The model reconstruction process was then formulated as a mixed-integer linear programming (MILP) optimization problem, where the objective was to maximize the inclusion of reactions supported by gene expression while maintaining the functionality of the essential tasks. Reactions without strong expression support were penalized but retained if required to complete any essential metabolic function. The constructed model was finally manually curated by first removing dead-end metabolites and reactions to streamline the metabolic models. The resulting curated model contained 4168 reactions and 2483 metabolites (**Supplementary Table 8**, provided as separate excel file).

#### 1.2. 13C-MFA flux simulation

In order to estimate the flux rate leveraging 13C-glucose isotopomers for DMSO, sertraline, carfilzomib and the combined treatment groups, we constructed the isotopomer model from the RPMI-8402 cell line-specific model. As the 13C-glucose tracing measured the isotopomers for the reactions in central carbon metabolism, including pathways like glycolysis, TCA cycle and pentose phosphate pathway, a smaller version of the metabolic model was developed that contained 200 reactions in central carbon metabolism, including 28 exchange reactions and biomass reaction.

We then constructed the Atom Mapping Matrix (AMM) to represent carbon atom transitions from substrates to products in the metabolic reactions using the MetAMDB database[5], which are essential for constructing Elementary Metabolite Units (EMUs). For reactions not available in MetAMDB, we employed the Reaction Decoder Tool (RDT) toolkit to compute AMM using reaction SMILES representations.

The measured isotopomer fractions were represented as Mass Distribution Vectors (MDVs), where each element corresponded to the relative abundance of a specific isotopomer. To simulate these MDVs and relate them to intracellular fluxes, the EMU framework was employed, leveraging the Freeflux framework[1]. The EMU method establishes a functional relationship between fluxes and MDVs by breaking down the metabolic network into EMUs, representing the minimal subsets of metabolite atoms needed to model isotopic labelling[6]. For each target EMU, the method identifies the smallest set of precursor EMUs that contribute to its labelling[7], guided by an EMU adjacency matrix, which encodes the connectivity between EMUs based on atom transitions defined by the Atom Mapping Matrix (AMM).

The label fractions were iteratively propagated from precursor EMUs to product EMUs, enabling the simulation of MDVs across the network. Under steady assumption, the fluxes were estimated by solving a nonlinear least squares optimisation problem to minimise the sum of squared differences between simulated and experimentally observed MDVs (The 13C MFA fluxes are provided in **Supplementary Table 8**, provided as separate excel file).

#### 1.3. Integrating 13C fluxes and proteomics into the RPMI-8402 metabolic model.

The estimated fluxes from 13C MFA were then integrated into the RPMI-8402 model to improve the treatment condition-specific flux simulation. In particular, the upper and lower bounds of the reactions in the central carbon metabolism were constrained with the flux rates estimated by 13C MFA as in the equation below.

$$lb = -\gamma \cdot flux_{13c}$$

$$ub = \gamma \cdot flux_{13c} ,$$

where $flux_{13c}$ is the flux rates estimated from 13C MFA, used to impose condition-specific constraints on the model, and $lb$ and $ub$ represents the lower and upper bounds of the reaction fluxes, respectively. $\gamma$ = 10 is the hyperparameter used to relax the upper and lower bounds and allow a flexible flux range.

Then the sample-specific proteomics data were integrated into the model, leveraging the GECKO framework, where each enzymatic reaction was constrained by its corresponding protein abundance [8]. For a metabolic reaction $v_i$, catalysed by an enzyme $E_i$, the flux is constrained as $v_i \leq K_{cat,i} \cdot [E_i]$, where $K_{cat,i}$ is the enzyme's turnover number derived from the BRENDA database [9] and $[E_i]$ is its measured abundance. The stoichiometric matrix ($S$), where rows represent the metabolites and the columns represent the reactions, was augmented with enzyme usage reactions and a global constraint, $\sum [E_i] \leq E_{total}$, enforcing the total enzyme pool based on the cell's protein budget. Then, the sample-specific fluxes were estimated using Flux Variability Analysis (FVA), which estimates the minimum and maximum flux rates at steady state [10,11], and used for metabolic enrichment analysis across conditions.
